# Addressing challenges in agentic retrieval of structured data from biomedical databases

**DOI:** 10.64898/2026.04.25.720782

**Authors:** Abhishek Halder, Manvendra Singh, Rohit Kesarwani, Bernadette Mathew, Namrata Bhattacharya, Om Chikhaliya, Dhruv Motwani, Sreeram Chandra Murthy Peela, Swarnava Samanta, Himanshu Nagdev, Piyush, Prateeksha Muddemmanavar, Mohammed Farooq, Gaurav Ahuja, Debarka Sengupta

## Abstract

Biomedical research increasingly relies on expert-curated databases to connect diseases, genes, variants, phenotypes, pathways and therapeutics. Incomplete or irreproducible retrieval can distort interpretation, mechanistic inference, variant assessment or therapeutic prioritization despite an unchanged evidence base. Agentic natural-language-to-SQL (NL2SQL) systems and Model Context Protocol-enabled agents can autonomously retrieve evidence from biomedical databases, including Open Targets and the Highly Confident Drug-Target Database, in response to natural-language questions. However, they may truncate large result sets, miss records when terminology differs from database vocabulary and return different outputs across runs. We systematically quantify these failures and introduce BioChirp, a goal-directed autonomous system for reliable retrieval from structured biomedical databases, built on interpretation-execution separation. A coordinated language-model layer interprets intent and resolves database-field assignments, after which entity resolution maps user terms to database terminology. A deterministic Steiner-tree planner and executor then retrieve matching records without further language-model involvement. Across curated biomedical databases spanning more than one million associations, BioChirp retrieved thousands of records for exhaustive queries, whereas baseline agents returned at most a few hundred or failed. Across 70 queries, five runs and three BioChirp database backends, BioChirp achieved a median cross-run Jaccard similarity of 1.0. Across 910 database-question evaluations derived from expert-written BioASQ questions and spanning ten databases, BioChirp retrieved at least one database record in 85.4% of cases and produced a fully correct answer in 51.1% of cases, compared with 44.6% and 17.4% for agentic NL2SQL. BioChirp is publicly available at https://biochirp.iiitd.edu.in.

## Introduction

Complete and reproducible outputs are important when artificial intelligence systems support scientific evidence retrieval, because incomplete or inconsistent results can influence downstream analyses. Biomedical research relies on structured associations among diseases, genes, variants, phenotypes, proteins, pathways, chemicals and therapeutics to support disease interpretation, mechanistic analysis, variant assessment and intervention prioritization. A researcher investigating a biomedical condition may need to integrate all relevant phenotype–gene, variant– disease, gene–pathway and treatment associations before deciding which hypotheses to pursue. A missed association can affect the resulting biological interpretation. Biomedical knowledge retrieval therefore provides a practical test of agentic AI reliability across disease interpretation, variant assessment, mechanistic analysis and therapeutic research.

Much of this evidence resides in expert-curated databases, but retrieving it often requires programming skills, knowledge of database schemas and familiarity with resource-specific query interfaces. Agentic interfaces can reduce this barrier by interpreting natural-language questions and executing retrieval workflows with limited human intervention^1–5^. This convenience is valuable only if the agent retrieves all relevant evidence and returns the same result when the question and database remain unchanged.

Current systems often provide database access through agentic NL2SQL pipelines or Model Context Protocol-enabled agents. Agentic NL2SQL systems use language models to generate SQL queries, whereas MCP-enabled agents select tools and coordinate retrieval across connected resources. Both approaches simplify access, but they can fail in three ways. They may truncate large result sets^6–8^, miss records when the user’s terminology differs from the database vocabulary^9–11^ or return different results across repeated runs^12–16^. For example, a query for Gleevec may return no match if the database stores the drug only as imatinib. Truncation creates a different problem because the system may return a plausible answer while silently excluding many valid associations. Users may therefore receive an apparently complete answer even when the system has left out relevant records.

Relational databases make these failures especially consequential because they organize information differently from the document collections commonly used in retrieval-augmented generation. Document retrieval usually ranks semantically similar passages and puts a limited number of top-ranked chunks in the model context. Relevant facts may also recur across overapping passages, so missing one chunk does not necessarily remove the information needed to answer the question^17^. Relational databases instead encode evidence as discrete records distributed across linked tables. To answer a single biological question, the system may need to interpret the database schema, resolve entity names, select the relevant tables, construct the required joins and retrieve thousands or millions of associations. Large result sets can exceed a language model’s context window and increase the cost of processing the retrieved records. In relational databases, each association represents a distinct piece of evidence, so no other record can compensate for one that is missed. If the system misses a drug–target or gene– disease association, the answer may incorrectly suggest that the database contains no such association.

Despite these risks, researchers usually evaluate agentic database systems by checking whether they produce correct final answers. These evaluations rarely measure whether the system retrieves all matching records or returns the same evidence across repeated runs. We therefore evaluate truncation of large result sets, failure to recognize alternative terminology and variation across repeated runs. Although we study these problems in biomedical databases, they are not unique to biomedicine. Agents that retrieve large, terminology-sensitive record sets from relational databases can face similar failures. When downstream decisions depend on the retrieved evidence, incompleteness and irreproducibility become safety and reliability risks because the same query can expose users to different evidence and support different downstream conclusions.

We developed BioChirp to address these limitations through interpretation–execution separation. BioChirp operates as a goal-directed autonomous system that interprets user intent, plans a retrieval strategy and executes it without further human intervention. Its coordinated interpretation layer runs multiple language-model components in parallel, with an orchestrator resolving disagreements in database-field assignments before entity resolution. A deterministic Steiner-tree planner^18^ selects the required tables and constructs the retrieval path, after which a deterministic executor retrieves the matching records without further language-model involvement. This design preserves natural-language access while making retrieval deterministic and auditable after interpretation and entity resolution have been fixed.

We evaluate BioChirp across eleven biomedical databases through controlled tests of retrieval completeness, synonym robustness and cross-run reproducibility, together with a comparison against agentic NL2SQL across ten relational databases. BioChirp produced reproducible results across deterministic backends and substantially improved both database-record retrieval and answer correctness across 910 database–question evaluations derived from expert-written BioASQ^19^ questions. To our knowledge, this study provides the first systematic quantification of association-level completeness loss in language-model-based agents operating over curated biomedical databases.

## Results

### Overview of BioChirp

Many agentic retrieval systems rely on language models across multiple stages of the retrieval process. This approach simplifies database access but can omit relevant records. A model may fail to recognize an alternative biomedical name, select the wrong database field or stop after retrieving only a subset of the matching evidence. Repeating the same question can also produce a different evidence set. We therefore designed BioChirp to use language models only where linguistic or biomedical interpretation was required, while keeping retrieval planning and execution deterministic.

BioChirp separates question interpretation from database retrieval (Fig. 1A and B). Language models identify the biomedical entities in a question, determine which database fields should constrain the search and specify which fields should be returned. These decisions form a fixed retrieval specification. Once the database fields have been finalized and the entity terms have been mapped to canonical database values, language model involvement in retrieval ends.

**Figure 1.**
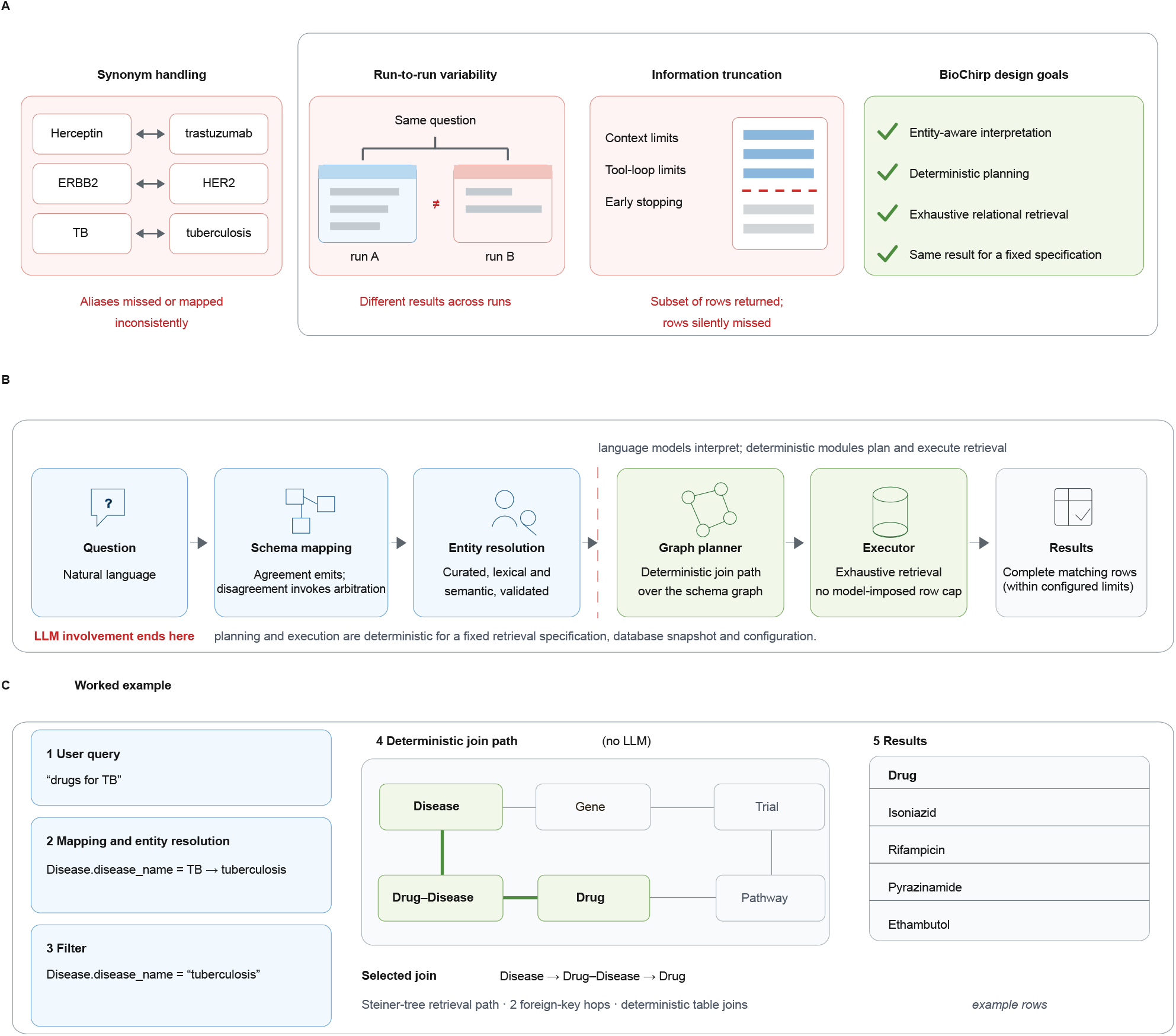
BioChirp separates language-model interpretation from deterministic retrieval. A, Three recurring limitations of language-model-mediated retrieval: synonym brittleness, runto-run variability and silent truncation of large result sets, together with the corresponding BioChirp design objectives. **B,** BioChirp workflow. A natural-language question undergoes coordinated schema mapping and entity resolution to produce a fixed structured retrieval specification. Agreement between two independent mappers emits the specification directly, whereas disagreement invokes orchestrator arbitration. Language-model involvement ends after interpretation; a deterministic schema-graph planner selects the required join path and the executor retrieves all records matching the fixed specification. **C,** Worked example for the query “drugs for TB”. The alias TB is resolved to tuberculosis, mapped to the disease-name filter and connected to the requested drug output through the deterministic Disease–Drug-Disease–Drug schema path. The resulting records are retrieved completely within the configured execution limits and remain identical for the same fixed specification and database snapshot.

Two rewriting models independently identify the biomedical entities in the question and clarify which entities constrain the search and which information should be returned. BioChirp uses each interpretation to identify a focused set of potentially relevant database fields. Two models independently determine which database fields should be filtered using the entities named in the question and which database fields should provide the requested answer. When both models select the same fields, BioChirp uses their assignments directly. When they select different fields, an orchestrator compares both outputs and determines which fields should be filtered and which should be returned.

BioChirp next maps each filter value to the terminology stored in the selected database. This step is required because users may describe a disease using an abbreviation, a drug using a brand name or a gene using a historical symbol. BioChirp uses synonyms obtained from authoritative biomedical databases to map alternative entity names to the canonical name or identifier used by the selected database. For example, Open Targets may provide evidence that HER2 and ERBB2 refer to the same gene, after which BioChirp uses the canonical value required by the target database. For free-text fields whose terminology varies across databases, BioChirp uses broader candidate-generation methods. It generates candidates through nomenclature expansion, approximate string matching and semantic similarity. A language model then removes candidates that do not refer to the intended entity. The accepted terms are converted to canonical database names or identifiers before retrieval begins.

For relational databases, BioChirp represents tables as nodes and inferred foreign-key relationships as edges in a schema graph. The selected filters and requested outputs identify the tables required to answer the question. A deterministic Steiner-tree planner connects these tables through the necessary association tables. The executor applies the filters to the appropriate database columns, performs the declared joins and returns all unique matching rows within the configured operational bounds. Open Targets uses predefined GraphQL operations instead of schema-graph planning. In both cases, the primary output is the structured evidence set. A language model summarizes the retrieved records, but it does not influence which records are included in the evidence set.

For the question “Which drugs are used to treat TB?”, BioChirp identifies TB as the disease constraint and drugs as the requested output. Entity resolution maps TB to tuberculosis. The planner then connects the Disease table to the Drug Disease association table and the Drug table. The executor applies the tuberculosis filter and returns the matching drug records (Fig. 1C). Repeating the query with the same resolved database field mappings, database snapshot and configuration returns the same set of records.

This design makes retrieval inspectable and reproducible after interpretation and entity resolution have been fixed. It does not guarantee that the original question was interpreted correctly. An entity that is missed or mapped incorrectly during interpretation cannot be recovered during deterministic execution.

We evaluated BioChirp through component-level tests, end-to-end retrieval benchmarks and multistep cross-database workflows (Fig. 2). These analyses assessed field mapping, entity resolution, retrieval completeness, reproducibility and cross-database coordination against agentic NL2SQL and MCP-based systems.

**Figure 2.**
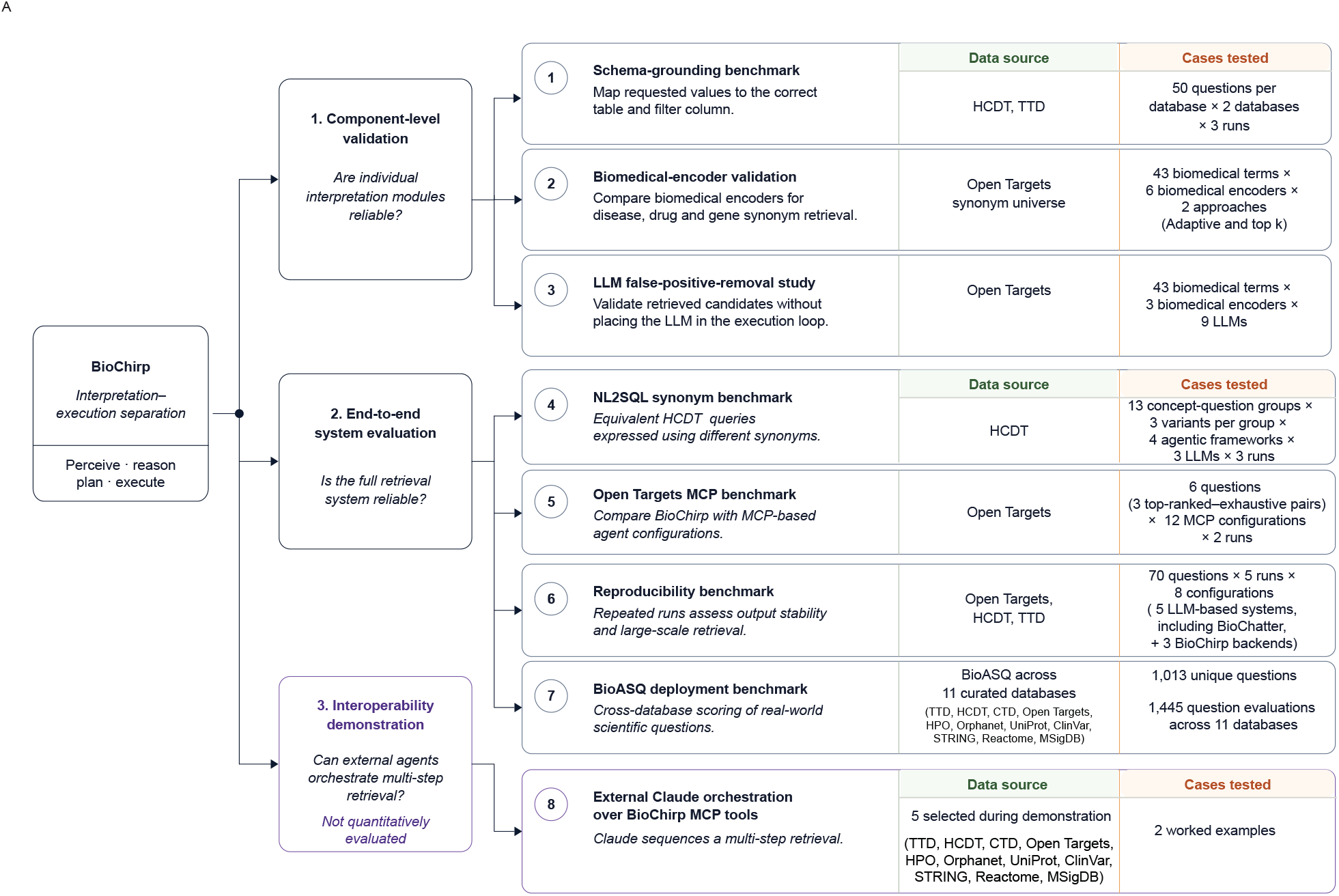
Evaluation framework for BioChirp component reliability, end-to-end retrieval and interoperability. The database-retrieval evaluation program comprised eight studies organized into three levels. Component-level validation included schema grounding on HCDT and TTD, biomedical-encoder evaluation using 43 terms and six encoders, and language-model filtering of candidates generated by three encoders. End-to-end evaluation included the HCDT synonym benchmark with 1,404 agentic NL2SQL runs, the Open Targets MCP benchmark with six questions and 12 configurations, a reproducibility benchmark with 70 questions, five runs and eight systems, and a BioASQ deployment study based on 1,013 unique questions. Because some questions were tested against more than one database, these produced 1,445 evaluations across the 11 curated databases, with each question counted once for every database in which it was tested (TTD, HCDT, CTD, Open Targets, HPO, Orphanet, UniProt, ClinVar, STRING, MSigDB and Reactome). Interoperability was demonstrated using two worked examples in which an external agent sequenced BioChirp retrieval tools across databases. These examples were not included as a quantitative benchmark. Model coordination was evaluated separately on MedQA independently of database retrieval. HCDT, Highly Confident Drug–Target Database; TTD, Therapeutic Target Database; MCP, Model Context Protocol.

### Tool-calling agents silently miss biomedical associations

Biomedical evidence queries often require complete disease–target, drug–disease or gene– disease association sets rather than a short list of leading results. We tested whether MCP-connected agents could retrieve large Open Targets association sets when explicitly asked for exhaustive retrieval. The benchmark included twelve configurations, with five using the OpenAI Agents SDK, five using the OpenAI chat endpoint and two using Claude Desktop (Methods; Fig. 3A; Supplementary Table 1). All configurations queried the official Open Targets MCP server. BioChirp answered the same questions through its non-MCP Open Targets pathway. The benchmark covered disease–target, drug–disease and gene–disease associations (Supplementary Table 2).

**Figure 3.**
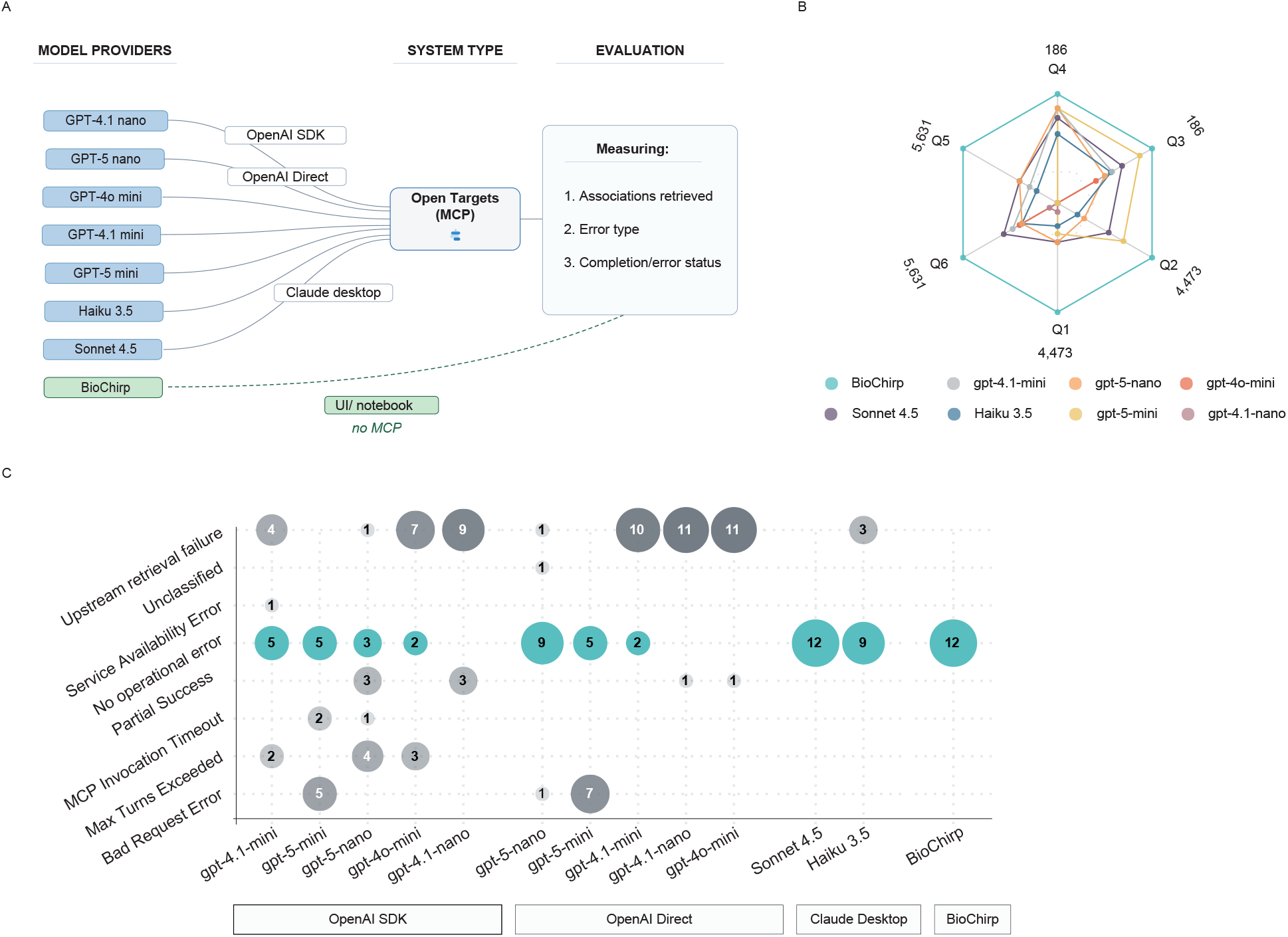
Incomplete association recovery across MCP-based agentic retrieval configurations. A, Benchmark configurations and measurements. Twelve combinations of OpenAI agentic-SDK, OpenAI direct-chat and Claude desktop-agent access were connected to the official Open Targets MCP server and compared with BioChirp retrieval outside the MCP loop. The number of retrieved associations, completion status and operational error type were recorded. **B,** Association count for six questions spanning disease–target, drug–disease and gene–disease associations in top-ranked and exhaustive settings. Values are shown relative to the corresponding BioChirp reference count, with the corresponding reference association counts indicated around the plot. The three top-ranked queries served as non-exhaustive operational controls and were not used to assess retrieval completeness. Count-yield comparisons were interpreted only for the three exhaustive queries. For the exhaustive chronic myeloid leukaemia query, Claude Sonnet 4.5, which completed all 12 benchmark runs without operational failure, returned 99 associations in both repetitions compared with a BioChirp reference count of 4,473. **C,** Distribution of completion and error outcomes across 12 runs per configuration (6 questions *×* 2 runs). Circle size and labels indicate the number of runs assigned to each outcome category. BioChirp completed all 12 runs successfully. [Model-label expansions: GPT-4.1 nano (gpt-4.1-nano); GPT-5 nano (gpt-5-nano); GPT-4o mini (gpt-4o-mini); GPT-4.1 mini (gpt-4.1-mini); GPT-5 mini (gpt-5-mini); Sonnet 4.5 (Claude Sonnet 4.5); Haiku 3.5 (Claude Haiku 3.5).] _31_

For each association type, one query requested the top-ranked associations and a paired query requested the complete set. For example, for chronic myeloid leukaemia, one query asked for the top-ranked molecular targets, whereas the paired query asked for all molecular targets associated with the disease. This design tested whether systems that handled a top-ranked retrieval task could also recover all matching associations.

The difference became clear when the queries required exhaustive retrieval (Fig. 3B). BioChirp answered both versions of the chronic myeloid leukaemia query and returned all 4,473 associated molecular targets. Among the MCP-connected configurations, Claude Sonnet 4.5 showed the most consistent successful operation and completed all twelve benchmark runs. However, it returned only 99 associations in both repetitions of the exhaustive chronic myeloid leukaemia query, corresponding to 2.2% of the BioChirp result count. Similar losses occurred for the exhaustive drug–disease and gene–disease queries. Other MCP-connected configurations also showed operational failures, including upstream retrieval errors, exceeded interaction limits, invocation timeouts and malformed requests. Some runs returned partial outcomes, whereas BioChirp completed all twelve runs without error (Fig. 3C). For count-based comparisons, each configuration was represented by the larger association count observed across its two runs, giving each MCP-connected system the benefit of its better observed retrieval outcome.

These findings distinguish successful tool calls from retrieval completeness. An agent could return a well-formed response while omitting most matching associations and provide no indication that further evidence remained. This benchmark compared association counts rather than record identities. A count below the exhaustive BioChirp reference establishes incomplete retrieval but does not determine which associations were omitted. Because the six questions were selected to stress large-result retrieval, the benchmark demonstrates that substantial truncation can occur under these conditions.

### Canonical names and synonyms produce inconsistent NL2SQL retrieval

Biomedical entities are often described using canonical names, brand names, abbreviations and historical aliases. Agentic NL2SQL systems must map these alternative forms to the values stored in a database before constructing an executable query. When an agent instead preserves the user’s wording in an exact-match filter, the query can return no records without producing an error. We tested whether this terminology-dependent failure persisted across agentic frameworks and language models despite explicit instructions to expand biomedical synonyms.

The benchmark included twelve NL2SQL configurations combining four agentic frameworks with three language models (Supplementary Table 3). Each configuration answered 39 questions arranged into 13 concept groups, with three terminology variants per group (Supplementary Table 4). The variants within each group asked the same retrieval question using different names for the same disease, drug or gene. For example, the imatinib group asked “Which genes does imatinib target?”, “Which genes does Gleevec target?” and “Which genes does STI571 target?”, using the drug’s canonical name, brand name and development code, respectively. The prompt explicitly instructed every configuration to identify known synonyms before constructing the query. Each question was repeated three times against the same HCDT snapshot, yielding 1,404 agentic NL2SQL runs (Fig. 4A). BioChirp answered the same 39 questions.

**Figure 4.**
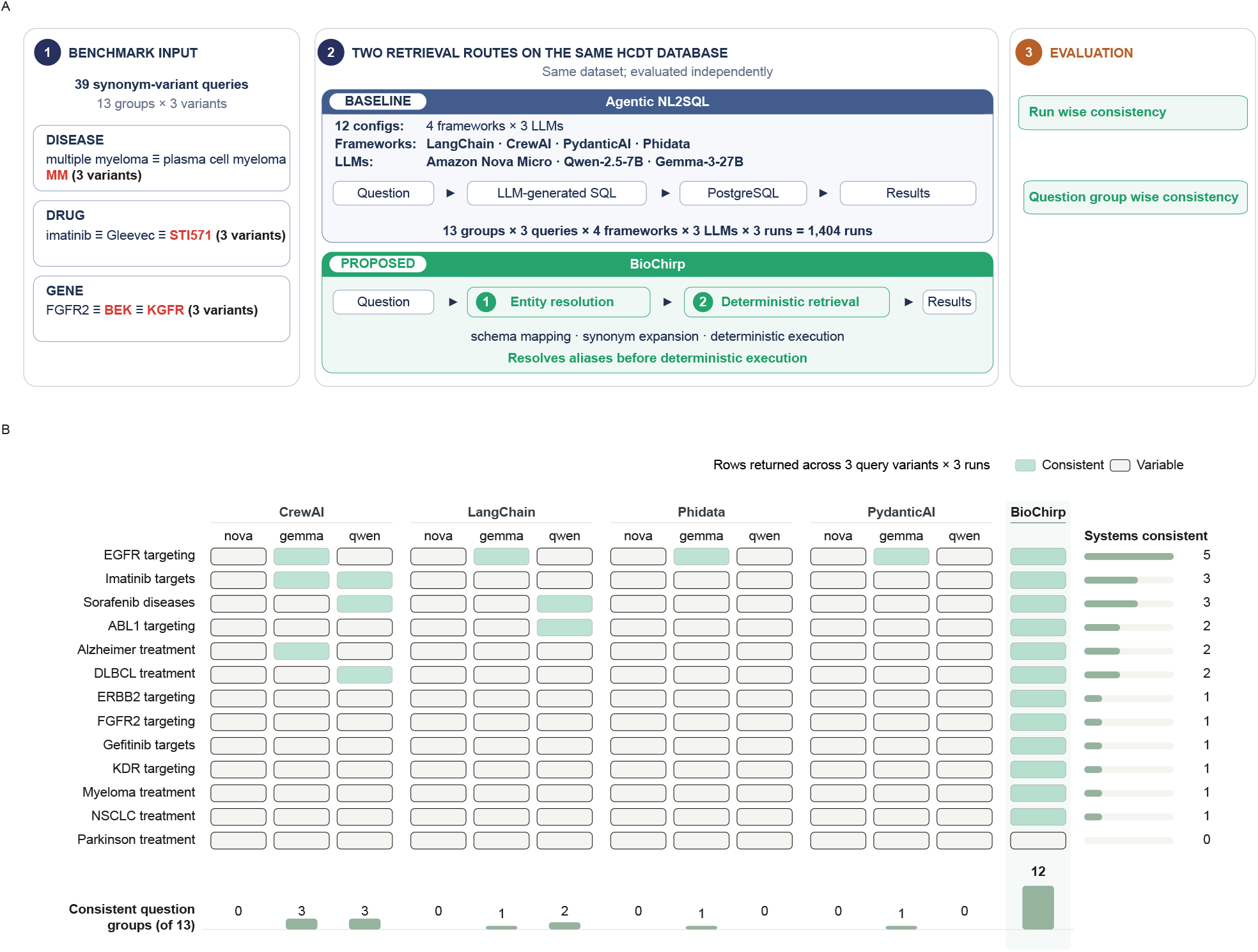
Agentic NL2SQL systems return inconsistent results across alternative biomedical names. A, Benchmark design. Thirty-nine HCDT questions were arranged into 13 concept groups, each containing three terminology variants referring to the same disease, drug or gene. Twelve agentic NL2SQL configurations, comprising four frameworks paired with three language models, generated and executed PostgreSQL queries. Each question was repeated three times, yielding 1,404 agentic NL2SQL runs. BioChirp resolved alternative entity names before deterministic retrieval from the same HCDT database. **B,** Stability of returned row counts across terminology variants and repeated runs. A configuration was considered stable for a concept group only when all three terminology variants returned the same row count in all three runs, giving nine executions per group. Green cells indicate groups meeting this criterion, whereas unfilled cells indicate variation. Marginal summaries show the number of configurations that were stable for each concept group and the number of stable groups for each configuration. BioChirp was stable for 12 of 13 groups. The strongest agentic NL2SQL configurations were stable for 3 of 13 groups, and six configurations were stable for none. [Model-label expansions: Qwen/Qwen-2.5-7B (qwen/qwen-2.5-7b-instruct); Gemma/Gemma-3-27B (google/gemma-3-27b-it); Amazon Nova Micro (amazon/nova-micro-v1).]

Despite explicit instructions to expand biomedical synonyms, the systems showed two recurring problems. First, non-canonical terms sometimes returned no records when the canonical term returned data, and different names for the same entity produced different row counts. Second, repeated executions of an unchanged question produced different row counts. We assessed both failures jointly (Fig. 4B). A concept group was classified as count-consistent only when all nine executions, comprising three terminology variants repeated three times, returned the same number of rows. BioChirp met this criterion for 12 of 13 groups. The strongest NL2SQL configurations by this measure, CrewAI with gemma-3-27b-it and CrewAI with qwen-2.5-7b-instruct, each met it for only 3 of 13 groups.

These results show that instructing the models to expand synonyms did not prevent terminology-dependent retrieval loss or run-to-run variation. The same biomedical entity could produce different row counts when expressed using a synonym or when the identical question was repeated, even though the query completed without error. This benchmark assessed row-count consistency rather than record-level identity agreement and was limited to 13 concept groups in one HCDT snapshot.

### Synonym variation limits biomedical retrieval completeness

Reliable retrieval begins by identifying the biomedical entity mentioned in the question and mapping it to the canonical name or identifier used by the database. Biomedical databases commonly index diseases, drugs and genes under canonical names or identifiers, whereas users may refer to the same entities through abbreviations, brand names or historical aliases. Because BioChirp separates interpretation from execution, errors at this stage propagate directly into the deterministic retrieval plan. A missed disease synonym excludes every linked drug, target and pathway record stored under the canonical disease, whereas an incorrect drug mapping retrieves evidence for a related but different molecule. Entity-resolution recall therefore sets the upper bound on downstream retrieval completeness.

We tested whether fixed candidate counts or fixed similarity thresholds could recover alternative biomedical names reliably. The benchmark comprised 43 entities sampled from standard classifications, including 14 diseases from ICD-11, 14 drugs from ATC and 15 genes from Gene Ontology molecular-function categories. Representative examples included type 1 diabetes and psoriasis among diseases, aspirin and methotrexate among drugs, and TP53 and EGFR among genes. Curated Open Targets synonym sets served as references^20^ (Fig. 5A). Reference-set size varied substantially within every entity type, from 14 to 96 synonyms for diseases, 15 to 114 for drugs and 8 to 66 for genes. For example, idiopathic pulmonary fibrosis had 14 accepted synonyms, whereas glaucoma had 96. At *k* = 50, retrieval could recover no more than 50 of the 96 glaucoma synonyms, while only 14 of the 50 candidates retrieved for idiopathic pulmonary fibrosis could match its reference set. Consistent with this mismatch, none of the six biomedical sentence encoders (Supplementary Table 5) reached 0.9 recall at *k* = 50 for any entity type. Achieving 0.9 recall required 75–300 candidates for diseases, 300–475 for drugs and 100–275 for genes (Fig. 5E–G). Retrieving a fixed number of candidate terms for every entity was unsuitable because entities with many valid synonyms required a larger candidate pool to maintain high recall, whereas applying the same large pool to entities with few synonyms introduced more false matches.

**Figure 5.**
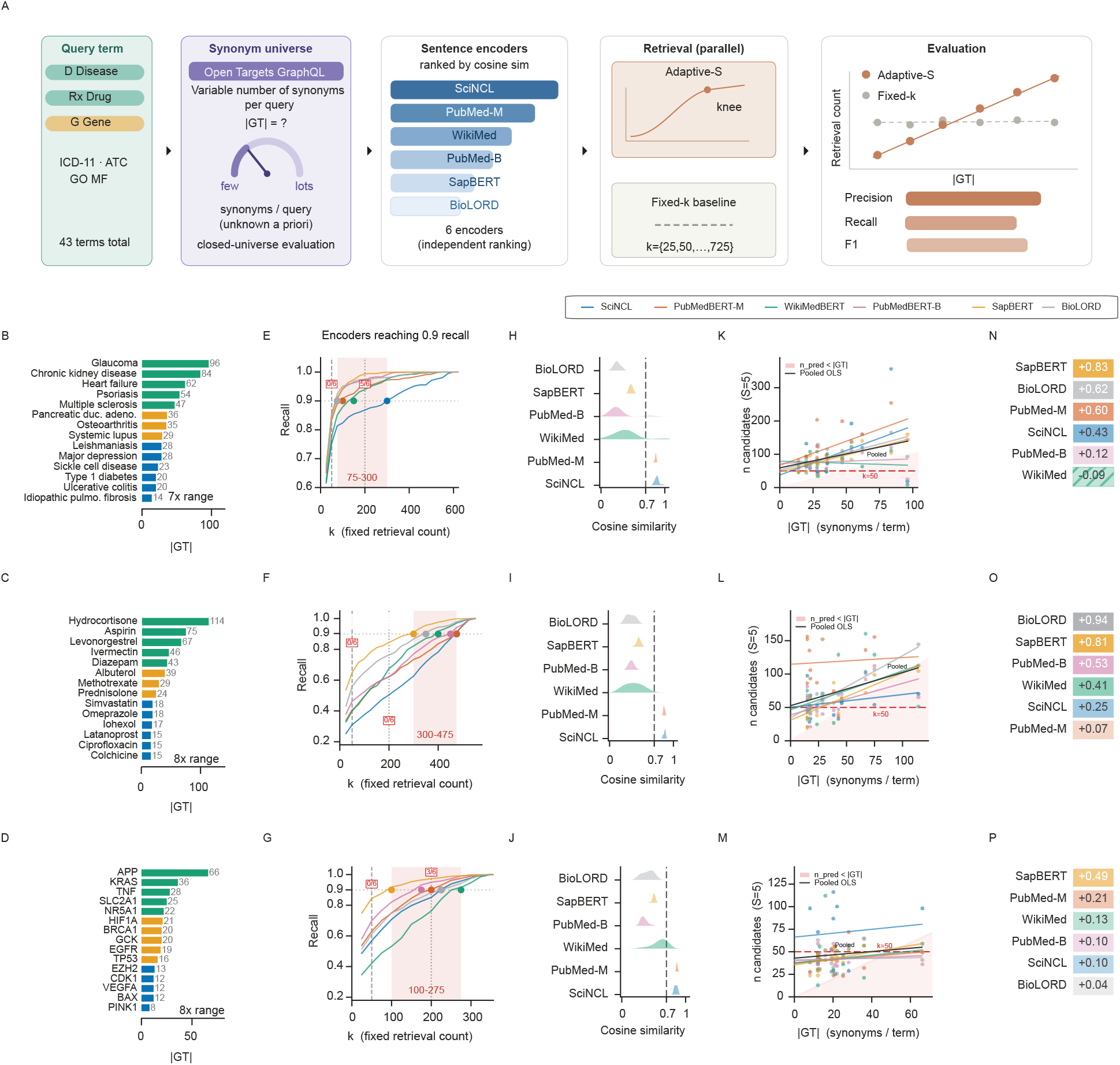
Query-adaptive candidate retrieval accommodates variation in biomedical synonym-set size. A, Evaluation design. Forty-three terms, comprising 14 diseases, 14 drugs and 15 genes sampled using ICD-11, ATC and Gene Ontology molecular-function categories, were evaluated against curated Open Targets synonym sets. Six biomedical sentence encoders independently ranked candidates by cosine similarity. A knee-based adaptive retrieval rule, Adaptive-S with *S* = 5, was compared with fixed candidate counts of *k* = 25, 50, . . . , 725. **B–D,** Number of reference synonyms, *|GT |*, for diseases, drugs and genes, respectively. Reference synonym counts vary seven- to eightfold within each entity type. **E–G,** Recall as a function of fixed retrieval count. Annotations indicate the candidate-count ranges required to reach 0.9 recall. No encoder reached 0.9 recall at *k* = 50 for any entity type. **H–J,** Distributions of cosine-similarity scores at the per-term Adaptive-S knee (*S* = 5) for each encoder, shown separately for diseases (**H**), drugs (**I**) and genes (**J**). The dashed line marks a fixed cosine-similarity threshold of 0.7. Variation across terms and encoders shows that a single fixed similarity cutoff does not generalize across embedding models. **K–M,** Adaptive candidate count as a function of reference synonym-set size for diseases, drugs and genes. Points represent individual terms and fitted trends summarize encoder-specific and pooled relationships. Under the knee cutoff, candidate count increases with synonym-set size rather than remaining fixed. **N–P,** Pearson correlations between adaptive candidate count and reference synonym-set size for diseases, drugs and genes. 33

We next examined whether a fixed similarity threshold could adjust retrieval depth to each entity. We estimated a query-specific cutoff from the knee of each ranked similarity curve using Adaptive-S with *S* = 5. Here, *S* is the Kneedle sensitivity parameter. Larger values make knee detection more conservative by requiring a longer flattening of the ranked curve before a knee is declared^21^. The similarity score at this cutoff varied across query terms and encoders and frequently differed from the representative fixed threshold of 0.7 (Fig. 5H–J). A universal threshold of 0.7 would therefore exclude potentially relevant candidates for some entities while retaining too many weakly related candidates for others. We consequently used the knee of each ranked similarity curve as a query-specific cutoff. Under this rule, the number of retrieved candidates varied with the size of the reference synonym set rather than remaining fixed (Fig. 5K–M). However, several encoders showed only a weak correspondence between synonym-set size and the number of candidates retrieved, particularly for genes (Fig. 5N–P). SapBERT^22^ showed the most balanced relationship across the three categories, with Pearson *r* = 0.83, 0.81 and 0.49 for diseases, drugs and genes, respectively. It also provided the most balanced precision–recall trade-off, with precision of 0.40–0.42 and recall of 0.94, 0.68 and 0.84 for diseases, drugs and genes, respectively (Supplementary Fig. 1D–F). Query-adaptive retrieval therefore accommodated entity-specific variation better than a universal candidate count or similarity threshold, but it did not eliminate false matches.

We therefore tested whether a language-model validation stage could remove these false matches without discarding too many valid synonyms. For diseases, SapBERT precision of 0.41 meant that 59% of retrieved candidates were not accepted synonyms. We evaluated nine language models (Supplementary Table 6) as zero-shot per-candidate filters over candidate pools generated by three encoders. In total, 1,161 encoder–model–term combinations were scored against the reference synonym sets (Fig. 6A). Every filter increased precision beyond the corresponding unfiltered encoder, but retained recall ranged from 0.27 to 0.74 across entity-type– language-model combinations (Fig. 6B–J). Language-model validation therefore removed many incorrect candidates, but it also rejected some accepted synonyms.

**Figure 6.**
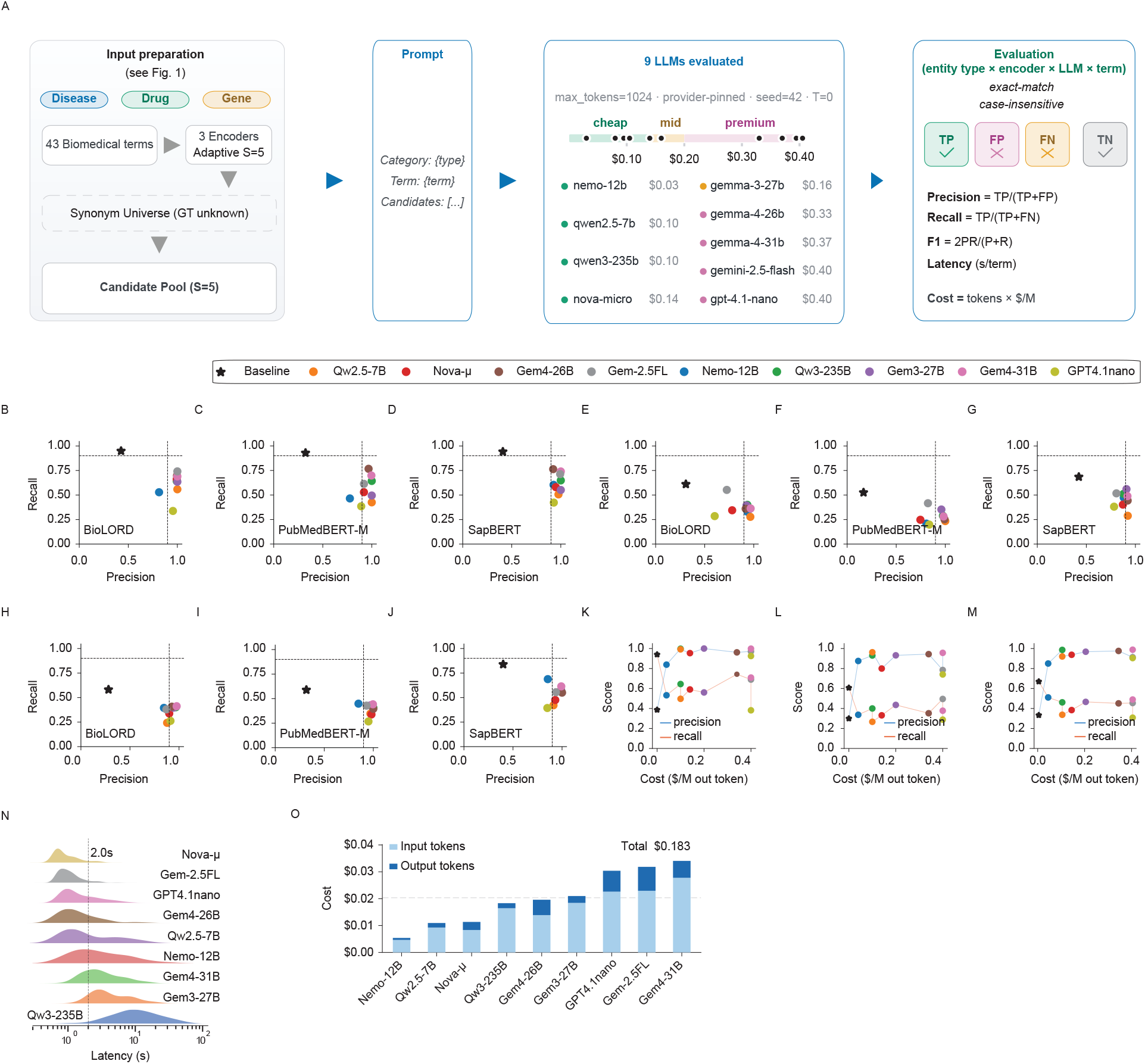
Language-model filtering increases candidate precision, but filtering quality is not ordered by model price. A, Benchmark design. Adaptive candidate pools generated for 43 biomedical terms by BioLORD, PubMedBERT-MARCO and SapBERT were passed independently to nine language models using the same binary synonym-validation prompt and fixed inference settings. Predictions were compared with curated Open Targets synonym sets using exact case-insensitive matching, yielding 1,161 encoder–model–term evaluations. **B–D,** Precision and recall for disease candidates generated by BioLORD, PubMedBERT-MARCO and SapBERT, respectively. **E–G,** Corresponding results for drug candidates. **H–J,** Corresponding results for gene candidates. Colored points represent language-model filters and black stars represent the unfiltered candidate pools. Filtering increased precision above 0.84 in every encoder and model combination, whereas retained recall varied among models. **K–M,** Precision and recall as functions of output-token price for diseases, drugs and genes, averaged across encoders. Higher price did not consistently predict better filtering performance. **N,** Distribution of per-term filtering latency for each model. **O,** Input- and output-token costs for the complete benchmark. The total cost across all nine models was US$0.183. [Modellabel expansions: Nemo-12B (mistralai/mistral-nemo); Qw2.5-7B (qwen/qwen-2.5-7b-instruct); Qw3-235B (qwen/qwen3-235b-a22b-2507); Nova-*µ* (amazon/nova-micro-v1); Gem3-27B (google/gemma-3-27b-it); Gem4-26B (google/ gemma-4-26b-a4b-it); Gem4-31B (google/ gemma-4-31b-it); Gem-2.5FL (google/ gemini-2 . 5-flash-lite); GPT4.1nano (openai/gpt-4.1-nano).] _34_

Language-model filtering quality did not improve monotonically with model price. Across a thirteenfold cost range, precision differed by only a few percentage points in most comparisons, recall showed no monotonic relationship with price and median latency ranged from 0.77 s to 9.9 s (Fig. 6K–N). The complete filtering benchmark cost US$0.183 (Fig. 6O). Models differed mainly in how broadly they retained candidates. Pairwise Jaccard agreement was lowest for drugs, ranging from 0.28 to 0.63, and post-filter recall was also lowest for drugs (Supplementary Fig. 2). Drug terminology was therefore the most difficult of the three entity types.

Together, these results defined BioChirp’s production entity-resolution policy. Based on the comparative precision–recall results (Fig. 6B–J), we selected Gemma-4-31B as the production entity-resolution filter (Supplementary Table 7). Drug and gene fields are resolved through exact matching against curated synonym vocabularies because semantic similarity can conflate related but distinct entities, such as aspirin and nitroaspirin. For free-text fields that cannot be mapped reliably through curated vocabularies alone, BioChirp combines curated synonym expansion, approximate string matching and adaptive SapBERT retrieval followed by language-model validation. Entity-resolution recall therefore remains the principal upper bound on retrieval completeness before deterministic planning begins.

### Deterministic retrieval improves reproducibility and evidence recovery

Autonomous retrieval systems should return the same evidence when a query is repeated without changes to the database contents or retrieval settings. In biomedical research, inconsistent or incomplete retrieval can change which drugs or targets are prioritized for further investigation. Such run-to-run variation is a reliability risk because the same evidence request can lead to different downstream decisions. Researchers obtain such evidence through two different routes. Language-model systems provide convenient natural-language access, whereas expert-curated relational databases store structured biomedical associations. Although these systems operate differently, both act as evidence sources when researchers use their outputs for downstream analysis. We therefore compared two sources of biomedical evidence. Five language-model baselines answered from their internal knowledge, whereas BioChirp retrieved associations from curated biomedical databases (Supplementary Table 8). This comparison measured the extent to which language-model answers captured associations available in curated biomedical databases. BioChirp accessed Open Targets through its live integration and TTD and HCDT through local preprocessed snapshots. Each system answered the same 70 natural-language queries across five independent runs (Supplementary Table 9; Fig. 7A). We quantified within-system reproducibility using cross-run Jaccard similarity, defined as the size of the intersection between association sets returned in repeated runs divided by the size of their union. A value of 1 indicates identical sets, whereas 0 indicates no shared associations.

**Figure 7.**
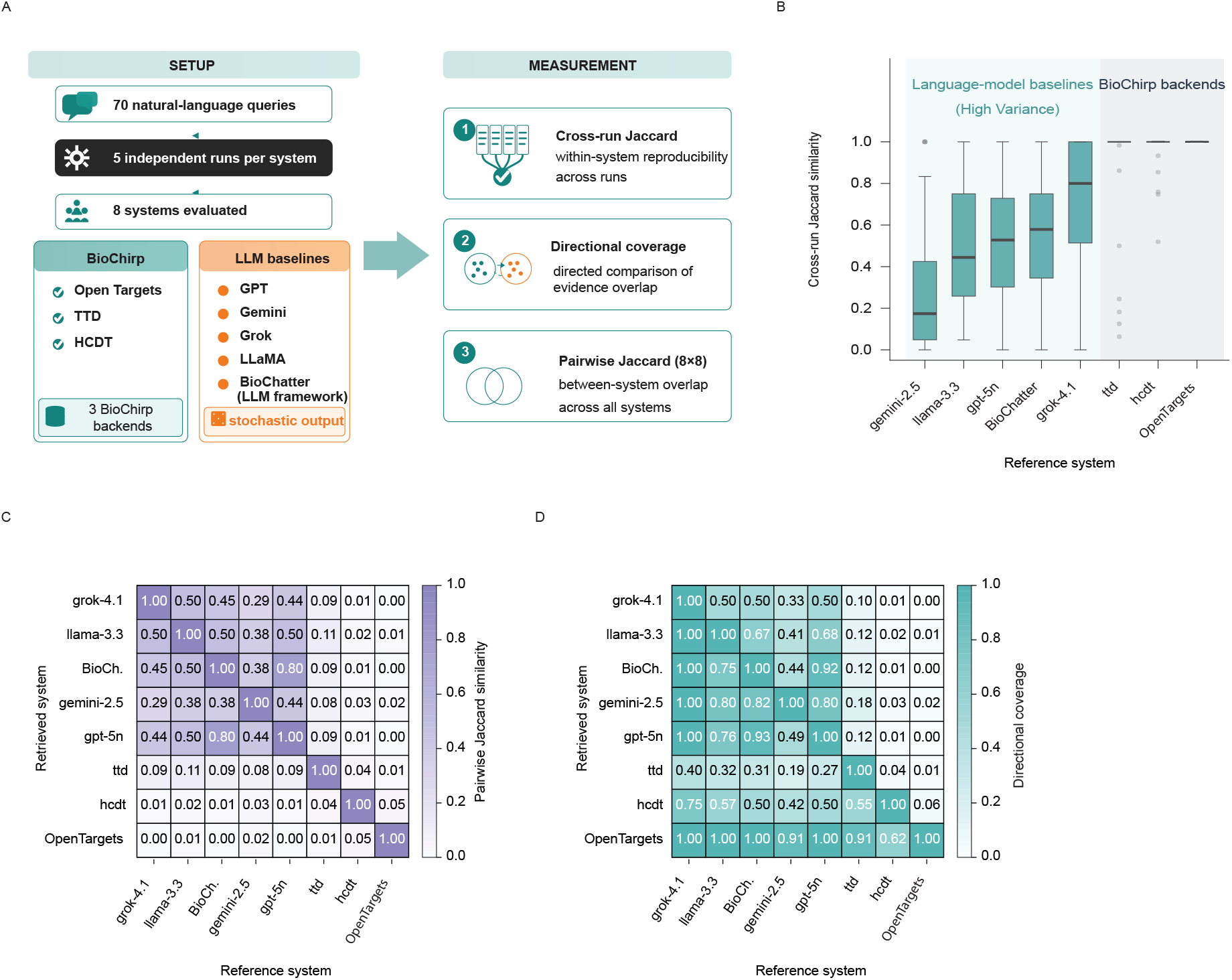
BioChirp improves repeated-query reproducibility and reveals asymmetric evidence recovery. A, Benchmark design. Five language-model baselines and three BioChirp database backends, Open Targets, TTD and HCDT, answered 70 natural-language questions over five independent runs. Reproducibility was measured using within-system cross-run Jaccard similarity, whereas between-system agreement was measured using symmetric Jaccard similarity and directional coverage. **B,** Query-level distributions of cross-run Jaccard similarity. The three backends achieved median Jaccard similarity of 1.0, whereas language-model systems showed lower and more variable reproducibility, with medians of 0.17 to 0.80. **C,** Symmetric pairwise Jaccard similarity between the unioned outputs of all systems. Agreement between any language model and any database backend did not exceed 0.11, whereas agreement between language-model pairs ranged from 0.29 to 0.80. **D,** Directed coverage matrix. For row system *i* and column reference system *j*, each cell reports *|S_i_ ∩ S_j_|/|S_j_|*, the fraction of the column system’s associations contained in the row system. Coverage was strongly asymmetric, particularly for Open Targets. Open Targets contained 91–100% of associations returned by language-model systems while those systems recovered 0–2% of Open Targets associations. [Model-label expansions: Grok/grok-4.1 (grok-4-1-fast-non-reasoning-latest); LLaMA/llama-3.3 (llama-3.3-70b-versatile); Gemini/gemini-2.5 (gemini-2.5-flash-lite); GPT/gpt-5n (gpt-5-nano); BioChatter (gpt-5-nano through the BioChatter wrapper).] _35_

BioChirp achieved a median cross-run Jaccard similarity of 1.0 across Open Targets, TTD and HCDT, indicating that the typical query returned identical association sets across repeated runs. By contrast, the five language-model baselines achieved median values of 0.17–0.80 and showed wide interquartile ranges (Fig. 7B). Among the language-model baselines, grok-4-1-fast-non-reasoning had the highest median cross-run Jaccard similarity. The language-model baselines nevertheless often produced different association sets across repeated runs of the same question. BioChirp was substantially more reproducible, but not uniformly identical across all evaluated queries.

Reproducibility alone does not show how much of the available evidence a system retrieves. We therefore quantified the proportion of associations in each curated database backend that every language-model baseline recovered. This analysis estimated the evidence loss that can occur when researchers rely on model-generated answers rather than direct database retrieval. We also measured overlap among the language-model outputs to determine whether different models repeatedly returned similar subsets. High overlap would be consistent with preferential recovery of associations that are more widely represented in model training data.

For each query, we treated the associations retrieved by BioChirp as the database result and measured how much of that result each language-model baseline recovered. Pairwise Jaccard similarity quantified the overall overlap between the model output and the database result. However, this value alone could not show whether a mismatch arose because the model missed associations present in the database or because it reported associations absent from the database result. We therefore calculated two complementary proportions. The first measured the proportion of model-reported associations also present in the database result. The second measured the proportion of database associations recovered by the model. These comparisons quantified retrieval coverage relative to the database contents and did not independently assess biological correctness.

Pairwise Jaccard similarity between any language-model output and the corresponding BioChirp database result did not exceed 0.11, whereas similarity among the language-model outputs ranged from 0.29 to 0.80 (Fig. 7C). The models therefore agreed more strongly with one another than with the associations retrieved from the databases. The directional analysis showed that this disagreement arose mainly because the models omitted database associations (Fig. 7D). For Open Targets, 91–100% of the model-reported associations were also present in the database result, yet the models recovered only 0–2% of the database associations. For HCDT, 42–75% of the model-reported associations were present in the database result, whereas the models recovered only 1–3% of the database associations. For TTD, the corresponding proportions were 19–40% and 10–18%. Across all three databases, many associations reported by the models were supported by the curated sources, but the models recovered only a small fraction of the associations available in databases. The language-model baselines therefore tended to converge on similar restricted evidence subsets. This pattern is consistent with preferential recovery of associations that are more widely represented in model training data, although the experiment did not test training-data exposure directly.

Together, these experiments revealed two reliability limitations of language-model-generated biomedical evidence. Language-model outputs varied across repeated runs and omitted large proportions of the associations available in curated databases. Either failure can alter downstream analyses and the biological conclusions drawn from them. By separating language-model interpretation from deterministic retrieval, BioChirp addresses these risks in biomedical applications where reproducibility and broad evidence coverage are safety-relevant requirements.

### Model coordination improves database field mapping

Before evaluating database field assignments, we tested whether coordinating multiple language models improved biomedical reasoning. We hypothesized that independently generated answers would provide complementary evidence that an orchestrator could compare and refine.

Across 100 MedQA questions and three runs, the coordinated system achieved a mean accuracy of 94.7%, compared with 92.0% for the strongest standalone model (Fig. 8A; Supplementary Note 1; Supplementary Fig. 3). The coordinated workflow outperformed every standalone model under the evaluated configuration.

**Figure 8.**
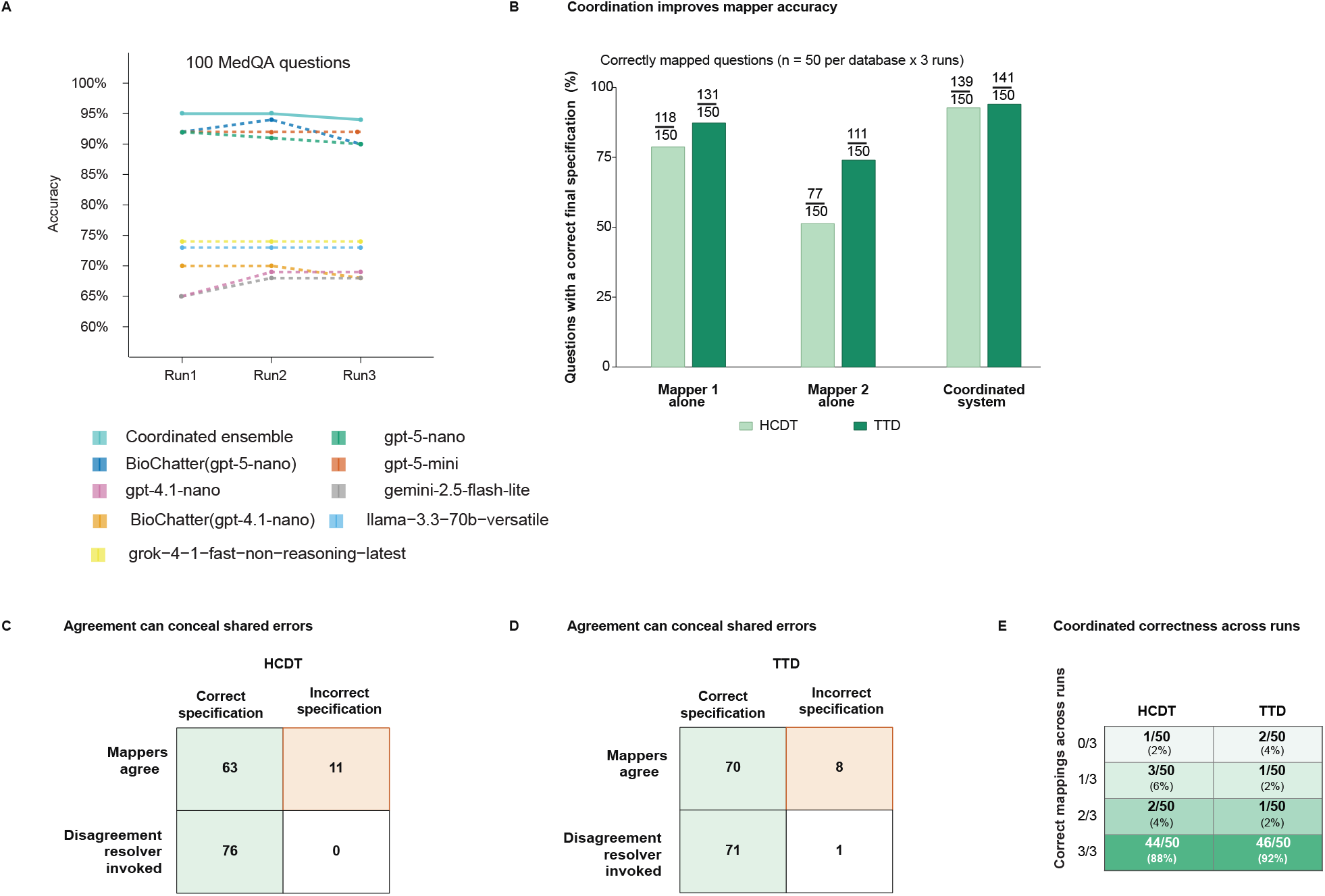
Multi-agent coordination improves biomedical reasoning and schema mapping, but agreement can conceal shared errors. A, Accuracy across three independent runs on 100 MedQA questions. Four heterogeneous language models answered in parallel and a fifth judge model reconciled their predictions. The coordinated ensemble achieved accuracies of 95%, 95% and 94%, exceeding all individual-model and BioChatter configurations. **B,** Schema-mapping performance on HCDT and TTD. Fifty questions per database were evaluated over three runs, giving 150 trials per database. Mapper 1, Mapper 2 and the coordinated system produced 118, 77 and 139 correct HCDT specifications and 131, 111 and 141 correct TTD specifications, respectively. **C,D,** Final specification correctness stratified by direct mapper agreement or invocation of the disagreement resolver for HCDT and TTD, respectively. Arbitration produced correct specifications for all 76 HCDT disagreements and 71 of 72 TTD disagreements. Eleven HCDT errors and eight TTD errors occurred when both mappers agreed on the same incorrect specification, which is the failure mode that agreement-gated coordination cannot detect. **E,** Number of correct final field mappings across three independent runs for each of 50 questions per database. The coordinated system produced a correct mapping in all three runs for 44 HCDT questions and 46 TTD questions.

We next tested whether model coordination improved how BioChirp translated biomedical questions into database field assignments. For each question, two models independently identified the biomedical entities, determined which database fields should be filtered and specified which fields should be returned as the answer. For example, in “Which diseases are treated with trastuzumab?”, the drug field should be filtered using trastuzumab and the disease field should be returned. When both models made the same assignments, BioChirp used them directly. When their assignments differed, an orchestrator examined both interpretations and produced the final field assignments (Supplementary Fig. 4A).

For each database, we developed 50 test questions and evaluated each question in three independent runs, giving 150 evaluations for HCDT and 150 for TTD. The first model independently produced the correct database field assignments in 78.7% of HCDT trials (118/150) and 87.3% of TTD trials (131/150). The second model correctly assigned the database fields in 51.3% of HCDT evaluations (77/150) and 74.0% of TTD evaluations (111/150). Combining the two interpretations through the orchestrator increased accuracy to 92.7% for HCDT (139/150) and 94.0% for TTD (141/150) (Fig. 8B).

We next examined how the orchestrator improved the final database field assignments. The two models disagreed in 76 of 150 HCDT evaluations and 72 of 150 TTD evaluations, causing these cases to be reviewed by the orchestrator. The orchestrator produced the correct final assignments in all 76 HCDT disagreements and in 71 of 72 TTD disagreements (Fig. 8C–D). In 10 of the 76 HCDT disagreements and 4 of the 72 TTD disagreements, neither model independently produced the complete correct field assignment. The orchestrator could therefore produce the correct database field assignment even when neither model produced it independently.

The remaining failures arose mainly when both models produced the same incorrect database field assignments. The models agreed in 74 of 150 HCDT evaluations and 78 of 150 TTD evaluations, but these outputs were incorrect in 11 HCDT cases and 8 TTD cases. Because agreement did not activate the orchestrator, the errors were retained in the final output. They accounted for all 11 remaining HCDT failures and 8 of the 9 remaining TTD failures. The orchestrator therefore corrected nearly all disagreements but could not detect errors made independently by both models.

The coordinated system produced a correct final field mapping in all three runs for 44 of 50 HCDT questions and 46 of 50 TTD questions (Fig. 8E). One HCDT question and two TTD questions were incorrect in all three runs, whereas the remaining five HCDT questions and two TTD questions were correct in only one or two runs. Overall, model coordination improved the accuracy of database field assignments and corrected nearly all disagreements, but it could not detect errors shared by both models or eliminate all run-to-run variation.

### BioChirp answers more biomedical questions using curated database evidence

Broad biomedical questions are difficult to evaluate against exhaustive answer keys because researchers may omit valid responses. For example, a manual search for biomarkers linked to diseases containing the word “cancer” would omit biomarkers linked to leukaemia, although leukaemia is a cancer. We therefore used BioASQ, which provides expert-curated biomedical questions and accepted reference answers. These references support evaluation against established answers without assuming that they enumerate every biologically valid association.

We compared BioChirp with an agentic NL2SQL baseline implemented using LangChain and qwen-2.5-7b-instruct (Fig. 9A). In the preceding terminology-variation benchmark, CrewAI with gemma-3-27b-it and CrewAI with qwen-2.5-7b-instruct returned identical record counts across all three terminology variants in 3 of 13 concept groups. LangChain with qwen-2.5-7b-instruct achieved this consistency in 2 of 13 groups. Initial runs showed that CrewAI was too slow for the full BioASQ evaluation. We therefore selected the LangChain–Qwen combination because it showed similar stability and could be evaluated at scale.

**Figure 9.**
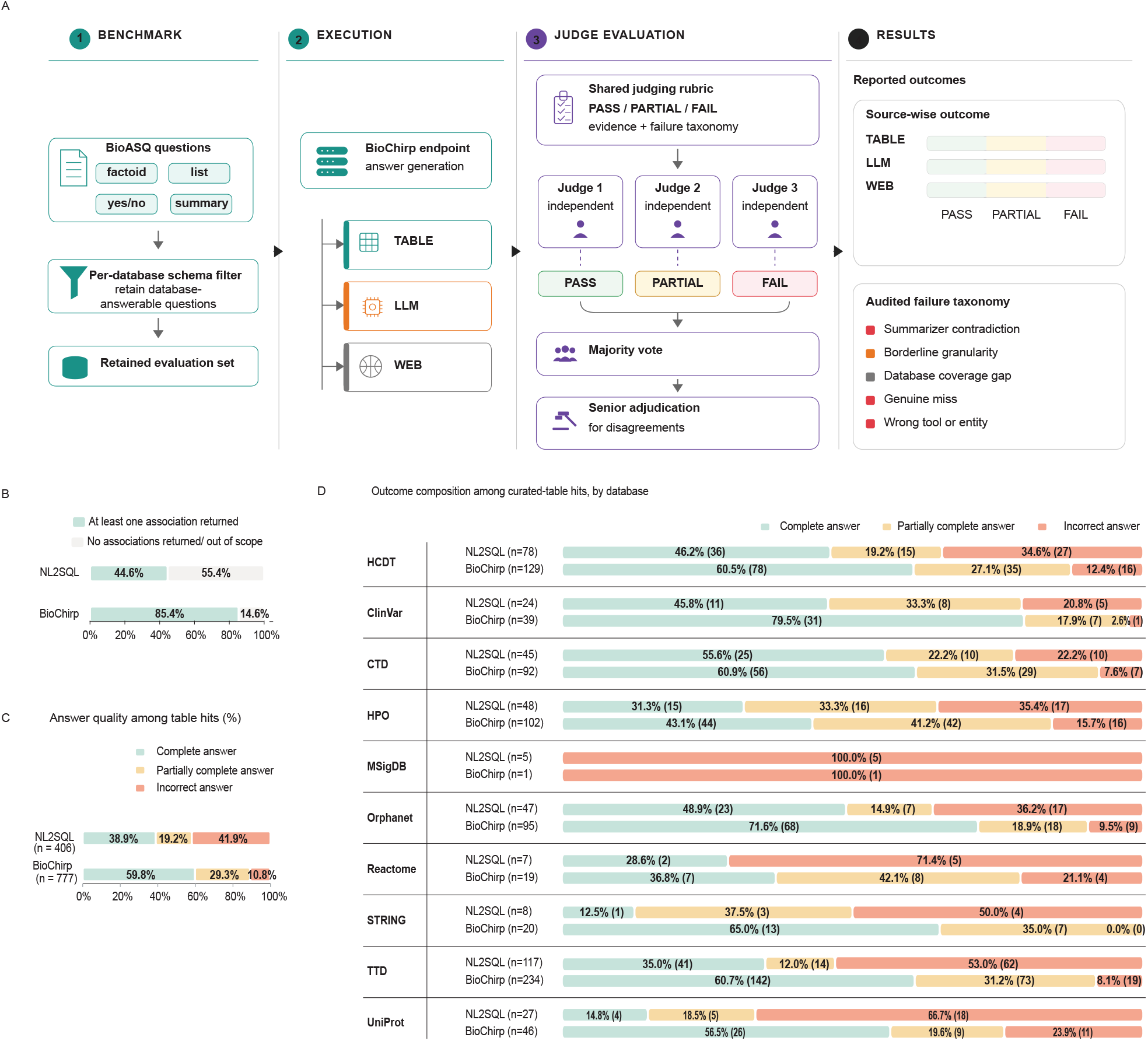
BioASQ-derived evaluation of BioChirp and comparison with agentic NL2SQL. A, Evaluation workflow. BioASQ factoid, yes/no, list and summary questions were screened against each database schema to retain those appearing answerable from the available content. BioChirp responses were grouped according to the source used to generate the answer, with TABLE indicating records retrieved from the assigned database, LLM indicating the language model’s internal knowledge and WEB indicating web search. Each response was independently evaluated three times by Claude Sonnet 5 using three predefined judge roles. Evaluations assigned complete, partially complete or incorrect labels, with the final verdict determined by majority vote. When all three labels differed, a fourth Claude Sonnet 5 evaluation using a senior-adjudicator role selected the verdict. Failure categories were audited separately. **B,** Proportion of 910 matched database–question evaluations with at least one association from the assigned database. BioChirp returned associations in 85.4% of evaluations, compared with 44.6% for agentic NL2SQL. **C,** Answer quality among table-hit evaluations. BioChirp produced more complete answers and fewer incorrect answers than agentic NL2SQL. **D,** Database-specific composition of complete, partially complete and incorrect answers among table hits for HCDT, ClinVar, CTD, HPO, MSigDB, Orphanet, Reactome, STRING, TTD and UniProt. Percentages were calculated within each system–database combination. Counts are shown in parentheses, and *n* denotes table-hit evaluations. _37_

The BioASQ source contained 5,729 unique questions spanning factoid, yes/no, summary and list formats. For each database separately, we used the ChatGPT web interface with GPT5.6 Thinking to review the database schema together with the complete BioASQ question set and identify questions that appeared answerable from that database. We retained the original wording of each question. A question could be assigned to more than one database, yielding 1,445 database–question evaluations from 1,013 unique questions across eleven databases.

Open Targets contributed 535 evaluations but was excluded from the agentic NL2SQL comparison because it uses a GraphQL interface rather than the relational schema required by the baseline. The matched comparison therefore contained 910 database–question evaluations across ten relational databases. BioChirp and agentic NL2SQL received the same question set within each database.

We assessed retrieval and answer quality separately. A table hit indicated that the system retrieved at least one association from the assigned curated database. When BioChirp retrieved no matching association, its predefined fallback pathway generated a response using either the language model’s internal knowledge or web search, and we classified the evaluation as having no table hit. All responses were evaluated against the BioASQ reference, but answer-quality percentages for the matched BioChirp–NL2SQL comparison were calculated only among table-hit responses. PASS indicated recovery of all expected reference elements, PARTIAL indicated recovery of a subset and FAIL indicated that the expected answer was not produced. Associations retrieved from the database but absent from the BioASQ reference were not automatically marked as incorrect.

Because manual review of all outputs across eleven databases would have been difficult to scale and standardize, we used a structured LLM-based evaluation workflow. Each answer was evaluated three times by Claude Sonnet 5 using 3 predefined judge profiles representing conservative factual assessment, biomedical domain-aware assessment and literal agreement with the benchmark reference. All profiles applied the same PASS, PARTIAL and FAIL criteria. A label selected by at least two judges became the final verdict. When the three evaluations returned different labels, a fourth Claude Sonnet 5 evaluation used a senior-adjudicator profile to select the verdict best supported by the retrieved evidence and reference answer.

Across the 910 matched evaluations, BioChirp returned at least one curated database association in 777 cases (85.4%), compared with 406 cases (44.6%) for agentic NL2SQL. When calculated across all 910 matched evaluations, PASS outcomes accounted for 51.1% (465/910) for BioChirp and 17.4% (158/910) for agentic NL2SQL. Among BioChirp’s 777 table-hit responses, 465 were classified as PASS (59.8%), 228 as PARTIAL (29.3%) and 84 as FAIL (10.8%). Among the 406 table-hit responses from agentic NL2SQL, 158 were classified as PASS (38.9%), 78 as PARTIAL (19.2%) and 170 as FAIL (41.9%). The remaining 133 BioChirp evaluations (14.6% of 910) and 504 agentic NL2SQL evaluations (55.4%) had no table hit and were excluded from these answer-quality percentages.

The table-hit advantage extended across the benchmark rather than arising from a single database. BioChirp returned curated database evidence for more questions than agentic NL2SQL in nine of the ten matched databases. MSigDB was the only exception. Few selected BioASQ questions aligned with the MSigDB schema, and neither system answered them correctly using database records. We therefore evaluated MSigDB separately with 4 additional schema-aligned questions (Supplementary Table 10). Although Open Targets was excluded from the matched comparison, BioChirp returned at least one curated database association for 470 of its 535 Open Targets evaluations, the largest single-database evaluation in the study.

We next re-audited all 157 BioChirp FAIL verdicts across the complete eleven-database evaluation, including TABLE, LLM and WEB answer pathways. Database coverage gaps accounted for 60 cases. In 29 genuine retrieval misses, the database contained the expected evidence but BioChirp failed to retrieve it. In 31 cases, BioChirp retrieved the correct evidence but contradicted, omitted or mis-selected it during final answer synthesis. Incorrect entity resolution or database routing caused 11 failures, and 26 cases reflected borderline evidence granularity. Incorrect outcomes therefore arose from both agent-level errors and limitations or ambiguities in the underlying databases. This audit included all answer pathways, whereas the 84 FAIL outcomes in the matched comparison were restricted to BioChirp responses with a table hit.

These results show that BioChirp outperformed the tested LangChain–Qwen baseline at both stages of agentic retrieval. It reached curated evidence more often and converted retrieved evidence into accepted answers more reliably. Its higher PARTIAL rate occurred alongside a markedly lower FAIL rate, indicating that incomplete BioChirp answers more often retained a valid subset of the reference rather than producing an incorrect answer. This pattern is consistent with interpretation–execution separation. BioChirp resolves terminology and identifies the relevant schema elements before deterministic planning and execution. The NL2SQL baseline instead relied on one language model to interpret terminology, navigate the full schema and generate the executable query.

The benchmark does not measure exhaustive biological completeness because BioASQ references may omit valid alternatives. The LLM-based question-to-database assignments were not exhaustively verified by human reviewers, and all judge profiles used the same underlying model. The experiment also did not isolate the individual contributions of terminology resolution, schema restriction and deterministic planning. The comparison therefore supports the complete BioChirp architecture against the tested agentic NL2SQL baseline rather than establishing superiority over every possible NL2SQL implementation.

### BioChirp supports browser and MCP-based agent access

To provide natural-language access across multiple biomedical databases, we deployed BioChirp as a live service at biochirp.iiitd.edu.in. The service provides access to eleven biomedical resources. Ten are served from local snapshots, whereas Open Targets is queried through a dedicated live API. Each resource is deployed as an independent backend service. The snapshot-backed services share microservices for schema mapping, entity resolution and deterministic retrieval planning, together with a common prompt template and sentence-embedding model. Database-specific schema descriptions, constraints, examples and deployment settings are supplied through configuration. This design allows individual database services to be updated or restarted independently.

Through the browser interface, users select a database-specific endpoint and submit a natural-language question. BioChirp returns the structured result together with the resolved filter terms, allowing users to inspect how the question was resolved without knowing the underlying schema (Supplementary Fig. 5). A separate MCP gateway exposes the same database endpoints to external agents.

The deployment allows both browser users and external agents to query the same BioChirp database endpoints, while each database service can be maintained and updated independently. It demonstrates interactive access and interoperability, but we did not evaluate uptime, throughput, load capacity or fault recovery. The quantitative evaluations bypassed the public browser interface and sent queries directly to the underlying database endpoints. They therefore assessed retrieval performance rather than the reliability or usability of the public web service.

### BioChirp MCP supports cross-database evidence integration

Complex scientific questions often require evidence distributed across multiple databases because no single resource captures all relevant biological relationships. An external agent must therefore identify the appropriate database for each part of the question, carry retrieved entities between databases and integrate the returned records into a coherent answer. We tested whether an external Claude agent could use BioChirp as a deterministic retrieval layer for this cross-database workflow.

Figure 10A illustrates this workflow with a question about RET that links its interaction partners to relevant pathways and associated chemicals across STRING, Reactome and CTD. Claude first queried STRING through BioChirp to identify proteins that physically interact with RET. It then used RET and selected interactors to retrieve related pathways from Reactome. Finally, it passed selected pathway-associated genes, including MAPK1 and MAPK3, to CTD^23^ to identify associated chemicals. This sequence connected molecular interactions, signalling pathways and compound associations. The CTD retrieval returned 769 associated chemicals before Claude generated the final database-grounded synthesis.

**Figure 10.**
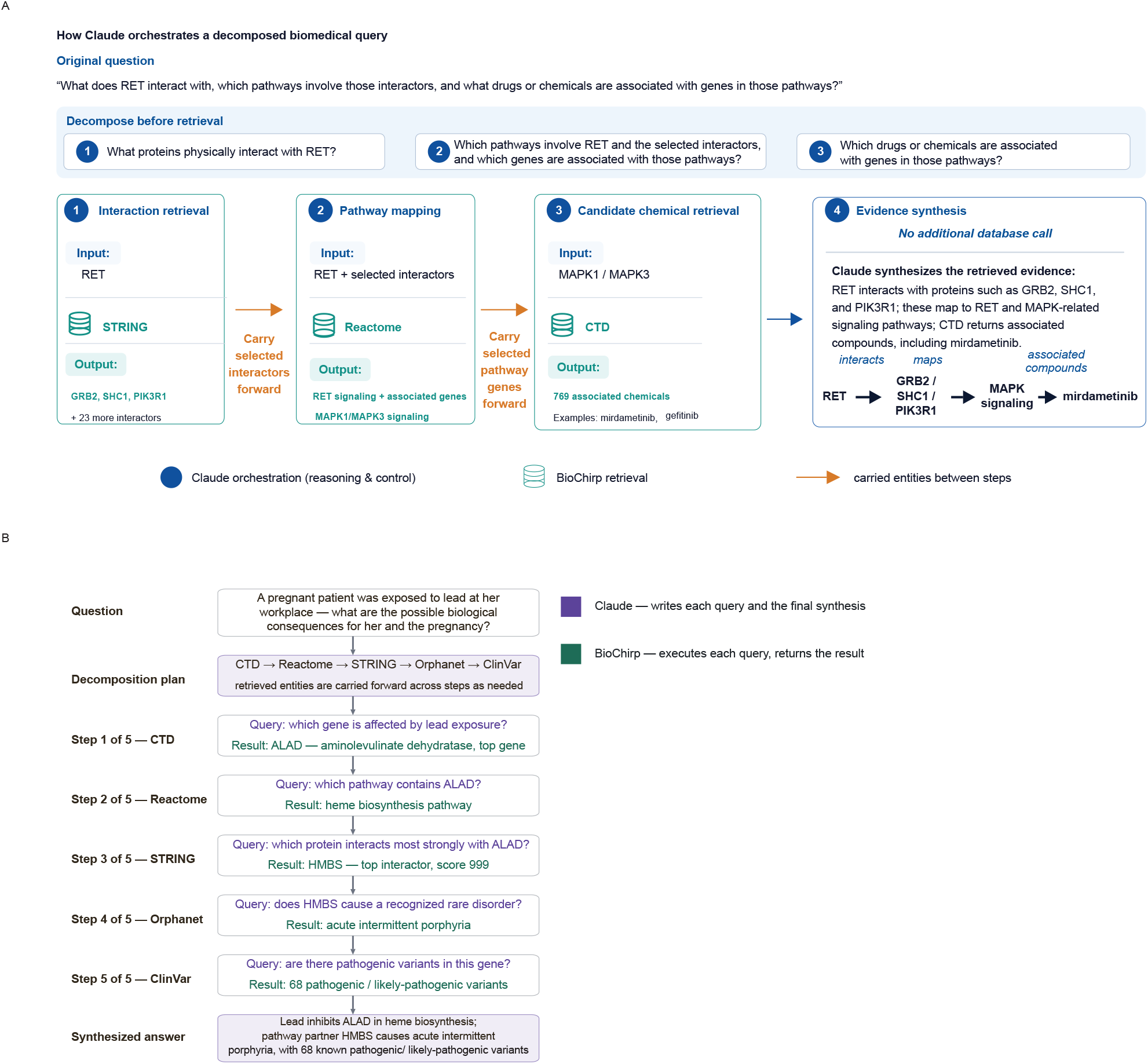
Claude-orchestrated decomposition and retrieval for a multistep biomedical question. A, Illustrative example showing how Claude decomposes a composite question about RET into sequential retrieval tasks executed by BioChirp. BioChirp first queries STRING and retrieves RET-interacting proteins, including GRB2, SHC1 and PIK3R1, together with 23 additional interactors. Claude carries selected interactors forward to Reactome to identify RETand MAPK1/MAPK3-related signalling pathways, and subsequently passes pathway genes to the Comparative Toxicogenomics Database (CTD), which returns 769 associated chemicals, including mirdametinib and gefitinib. Claude then synthesizes the retrieved evidence into a final response without an additional database call. Blue markers denote Claude-based reasoning and control, green database symbols denote BioChirp retrieval, and orange arrows indicate entities transferred between retrieval steps. **B,** A second illustrative example in which Claude decomposes a question about lead exposure into five sequential queries across CTD, Reactome, STRING, Orphanet and ClinVar. Claude uses the retrieved entities to formulate subsequent queries and integrates the evidence into the final answer. Purple denotes Claude-generated queries and synthesis, whereas green denotes results retrieved by BioChirp.

Figure 10B illustrates a second workflow that follows a question about lead exposure during pregnancy from an affected gene to a related pathway, protein interactor, rare disease and clinically interpreted variants across five databases. Claude carried entities sequentially across five BioChirp retrieval calls to CTD, Reactome, STRING, Orphanet^24^ and ClinVar^25^. The resulting chain linked exposure records, molecular pathways, protein interactions, rare-disease associations and clinically interpreted variants before the final answer was synthesized. This example represents an exploratory cross-database evidence chain.

In both workflows, Claude decomposed the original question, selected the database sequence, formulated each successive query, chose which retrieved entities to carry forward and synthesized the final response. BioChirp executed each database-specific query and returned the corresponding records.

The reliability of the final synthesis therefore remains under the control of the external agent. BioChirp only answers each decomposed database-specific query by retrieving the matching records. It does not decide which database to query next, which retrieved entities to carry forward or which evidence to include in the final response. Completeness and reproducibility therefore apply only to each individual BioChirp retrieval call, not to the overall cross-database workflow or final answer.

## Discussion

Natural-language interfaces allow researchers to retrieve evidence from biomedical databases without needing to understand their schemas, query languages or database-specific interfaces. This access is useful only when the system retrieves the relevant associations completely and consistently. A system that omits records or returns different evidence for the same request can mislead downstream analyses even when its final answer appears correct.

Methods for natural-language database access have advanced rapidly, but retrieval reliability has received less attention. Agentic NL2SQL systems generate executable queries, whereas tool-calling agents select functions and determine whether additional records should be retrieved. These systems are commonly evaluated according to the correctness of the final answer. Such evaluations do not establish whether all matching database records contributed to that answer. Our results show that silent truncation, terminology mismatch and run-to-run variation can remove relevant evidence even when the final response is judged correct. Retrieval completeness and reproducibility should therefore be evaluated separately from final-answer correctness.

BioChirp addresses these failures by restricting language model involvement to tasks that require linguistic or biomedical judgement. Language models interpret which information should constrain the search, what information should be returned and whether candidate entity names match the intended concept. Once the database fields and entity mappings are finalized, planning and record selection proceed deterministically. This separation prevents subsequent language model decisions from altering the retrieved evidence and therefore reduces run-to-run variation caused by query generation, join selection or iterative tool use. The configured operational limit sets an upper cap on the number of associations returned for a single request and reflects practical infrastructure constraints rather than a limitation of the deterministic retrieval design.

This separation also changes what BioChirp treats as its primary output. BioChirp returns structured relational records rather than only a natural-language summary, preserving the retrieved associations for inspection, filtering and downstream analysis. The finalized entity mappings and retrieval path also show how the original question was interpreted. Researchers can therefore examine the evidence independently of how it is summarized. A language model may present the records in readable form, but it does not determine which records are included in the evidence set.

The evaluated agentic systems retain language models within the database retrieval process. A model may generate the query, select database operations or decide whether additional records should be retrieved. Each of these decisions can alter which records enter the final evidence set. In BioChirp, language model involvement ends once the database fields and entity mappings have been finalized. Table selection, join-path construction and retrieval of matching records then proceed without further language model decisions.

Related studies have advanced language-model-based planning, retrieval and scientific tool use. The Modular Agentic Planner distributes planning across specialized language model modules^1^. Hyper-RAG uses a hypergraph to connect related medical information during retrieval^2^. iDesignGPT^26^ coordinates agents for engineering design, SciToolAgent^5^ uses a scientific tool knowledge graph to guide planning and tool use, and BioChatter^4^ combines language models with biomedical knowledge graphs and retrieval components. These systems address different tasks, but they retain language models in planning, tool selection or retrieval. BioChirp instead restricts language model involvement to question interpretation and entity mapping before database planning begins. The reliability of the subsequent retrieval therefore depends primarily on the quality of this initial interpretation, the coverage of the selected database and the configured operational limits.

Errors introduced during question interpretation, database-field assignment and synonym mapping to database vocabulary remain the main source of uncertainty in BioChirp. An incorrectly assigned database field or a missed synonym can exclude relevant associations even when planning and execution are correct. Biomedical terminology makes this stage difficult because multiple names may refer to the same entity, whereas lexically or semantically related terms may refer to different drugs, diseases, genes or biological processes. This difficulty was particularly evident for drug-name resolution, for which broader semantic candidate generation recovered fewer reference synonyms than it did for diseases and genes. Curated synonym resources therefore remain preferable when established alternatives are available, whereas broader lexical and semantic searches require careful candidate filtering.

Model coordination reduced errors in database-field assignment, particularly when the two models produced different interpretations of the same question. The orchestrator could compare these interpretations and recover the correct assignment in most disagreement cases. Coordination could not, however, detect cases in which both models made the same error. Agreement between models therefore indicates consistency but does not establish correctness. Future systems could add checks to confirm that the selected entities and database fields are valid and flag uncertain interpretations before retrieval begins.

BioChirp reduces incomplete and variable retrieval, but it cannot eliminate every failure. Synonym normalization reduces terminology mismatch, model coordination reduces field-assignment errors and deterministic execution removes variation from table selection, join construction and record retrieval. However, an incorrect interpretation, missed synonym or incorrect entity mapping can still exclude relevant records. BioChirp’s completeness claim therefore applies only after the database fields and entity mappings have been finalized. Within the configured upper limit, it retrieves all records matching that finalized request.

The retrieval process remains auditable. Users can inspect the finalized field assignments, entity mappings, selected tables, join path and returned records. When the interpretation deviates from the intended question, these intermediate outputs help identify the source of the error and allow the question to be clarified or paraphrased. The completeness claim applies to individual database calls rather than a cross-database reasoning chain with external agents. Reproducibility also depends on the database version, terminology-service outputs, language-model decisions and system configuration. BioChirp reduces variation at these stages where possible, but it cannot guarantee identical results when these inputs change.

Adding a new database to BioChirp requires its schema, field descriptions, terminology mappings and example questions to be prepared. This requires more setup than giving a language model direct database access. However, it makes the connection between the question and the returned records explicit. BioChirp is therefore most suitable for structured databases and tasks that require retrieval of all matching evidence.

BioChirp shows that language models do not need to control the full retrieval process. They can interpret the question, while deterministic components select tables, construct joins and retrieve records. This design makes missing evidence easier to detect and makes the limits of completeness and reproducibility clearer.

## Methods

### System overview

BioChirp separates natural-language interpretation from database planning and execution. A natural-language question *q* is submitted to a target database endpoint and converted into a structured retrieval specification. This specification records which database should be queried, which columns and values should constrain the search, and which columns should be returned. We denote it as *S*(*q*) = (*d, F, O*), where *d* is the selected database, *F* contains the resolved column–value filters and *O* contains the requested output columns.

Language models identify the biomedical entities in the question, determine which database columns should be filtered and returned, and validate alternative entity names during normalization to the database vocabulary. Once the database columns and entity values have been finalized, table selection, join construction and record retrieval proceed deterministically.

For relational databases, BioChirp identifies the required tables from the database schema and connects them through inferred foreign-key relationships. It then applies the finalized filters, performs the required joins and returns all unique matching records within the configured operational limits. For Open Targets, BioChirp maps the finalized request to a predefined GraphQL operation, supplies the resolved entity identifiers and retrieves all available pages within the configured operational limits.

In both cases, the retrieved database records constitute the primary output. BioChirp stores these records together with the applied filters, selected tables or operation, retrieval path and total result count, allowing the retrieval process to be inspected and audited. A language model receives the result statistics and a bounded preview of the retrieved rows only to generate a readable summary. It does not determine which records are included in the primary result set. The end-to-end workflow is summarized in Supplementary Algorithm 1.

### Data preprocessing

BioChirp accesses databases through local snapshots or live application interfaces. Open Targets^20^ is queried at runtime, whereas the other supported resources are processed locally before retrieval. Because the source formats and schemas differ across databases, preprocessing is database-specific. Here, CTD^23^, HCDT^27^ and TTD^28^ are used as representative examples of the preprocessing procedures rather than as an exhaustive description of all databases or columns. Across these examples, selected identifier and name columns were standardized, unnecessary prefixes and placeholder values were removed, and cells containing multiple values were split where required so that each resulting row represented a single entity or relationship (Supplementary Fig. 6A). When several identifiers were available for an entity, a fixed priority rule was used to select one canonical identifier (Supplementary Note 2).

The cleaned records were then organized into master tables containing canonical entity identifiers and names and association tables containing identifiers for related entities (Supplementary Fig. 6B). TTD preprocessing additionally retained biomarker records, mechanism-of-action categories and approval-status categories (Supplementary Fig. 6C). Duplicate rows, records with missing identifiers and malformed entries were removed. The complete preprocessing workflow is summarized in Supplementary Algorithm 2. Database-specific preprocessing workflows are available in the BioChirp GitHub repository (see Code availability).

### Language-model coordination for database field mapping

Biomedical databases can contain many tables and columns, although only a small subset is relevant to a given question. BioChirp therefore selects potentially relevant columns before assigning question elements to database fields, rather than presenting the complete schema to the language models (Supplementary Fig. 4A). Each database endpoint is configured with a curated set of queryable columns, which are the only fields that can be assigned as search constraints or requested outputs, together with rules for assigning question elements to those columns and example questions (Supplementary Note 3).

Before processing questions, BioChirp converts the database schema into vector representations for similarity-based column retrieval. Descriptions of database, table and column nodes are embedded using BAAI/bge-small-en-v1.5 (384 dimensions). For each queryable column, its embedding is combined with information from neighbouring nodes in the schema graph and then normalized. The resulting column vectors are stored in an in-memory Qdrant index and used to retrieve columns that are semantically similar to the rewritten question.

At query time, two language models independently rewrite the question to make the filter values and requested outputs explicit. Each rewrite is embedded with the same embedding model and used to retrieve the most similar queryable columns from the schema index. A language model reviews the columns retrieved for each rewrite and removes columns that are not relevant to the question. The retained columns from the two rewrites are then combined into a single candidate schema.

Two independent language models receive the same candidate schema and assign each filter value and requested output to a database column. Their mappings of filter values and requested outputs to database columns are compared programmatically. If both models produce the same mappings, they are accepted directly. If they disagree, an orchestrator reviews the two interpretations and produces the final field assignments. These assignments are then passed to entity resolution and subsequent retrieval planning. The complete field-mapping procedure is summarized in Supplementary Algorithm 3.

### Mapping biomedical terms to database vocabulary

After BioChirp determines which database columns should be filtered, the user-supplied values may still differ from the terminology stored in those columns. BioChirp therefore maps each filter value to the names or identifiers used by the selected database before retrieval planning begins (Fig. 1A).

A field-type router determines how each value is resolved. Database identifiers and predefined coded values, such as ICD codes, are matched directly to values stored in the database. Structured chemical representations, such as canonical SMILES strings, are also matched directly when stored in a standardized form. For gene and drug fields, BioChirp maps alternative names using synonyms and identifiers obtained from the corresponding database, its official API or established biomedical resources (Supplementary Note 4). For example, an official resource can map *Gleevec* to imatinib or *HER2* to *ERBB2*.

Disease names use a broader resolution procedure. For disease names, BioChirp combines synonyms obtained from databases, official APIs or established biomedical resources with lexical matching (Supplementary Note 5) and SapBERT-based semantic retrieval (Supplementary Note 6). For other free-text fields, including pathway names, BioChirp uses lexical and semantic retrieval. Lexical matching identifies variations caused by spelling, abbreviations or formatting.

Candidates produced by fuzzy or semantic retrieval are evaluated individually by a zero-shot language-model validator to remove terms that do not refer to the intended entity (Supplementary Note 7). Names obtained directly from the database, its official API or established biomedical resources bypass this validation step. The directly accepted names and validated candidates are combined, mapped to the corresponding database names or identifiers and deduplicated. These resolved names or identifiers become the final filter values used for database planning and retrieval. The complete entity-grounding procedure is summarized in Supplementary Algorithm 4.

### Deterministic retrieval planning

After BioChirp has determined which database columns should constrain the search, resolved the corresponding entity values to the database vocabulary, and identified which columns should be returned, it constructs the retrieval plan without further language-model involvement. Each relational database is represented as an undirected graph in which nodes are tables and edges correspond to inferred foreign-key relationships between tables.

The filter and output columns are first mapped to the tables that contain them. These tables define the terminals that the planner attempts to connect. BioChirp then uses a deterministic greedy Steiner-tree procedure to connect the terminals through the schema graph. Starting from one terminal, the planner repeatedly connects the nearest remaining terminal using the shortest available join path. Intermediate tables, including association tables not explicitly referenced in the query, are added when required to complete the connection (Supplementary Fig. 4C).

The resulting plan records the selected tables, join order and join-key pairs and is passed to the execution layer, which applies the filters and performs the specified joins. For a fixed retrieval specification and unchanged database schema, the same deterministic planning rules produce the same join plan.

### Retrieving records from relational databases

The executor applies the resolved filters, joins tables through the relationships specified by the plan and returns the requested columns. Duplicate output rows are removed.

Execution continues until all records matching the finalized filters and requested output fields have been processed or the configured operational limit has been reached. The operational limit is fixed before execution and reflects available memory, compute, storage, network capacity and endpoint pagination constraints.

Repeated successful executions return the same structured result when the finalized filters, requested output fields and database state remain unchanged. All databases except Open Targets were evaluated using local snapshots. Open Targets was queried through a live GraphQL endpoint and could therefore change during or after the evaluation period. The relational execution workflow is provided in Supplementary Algorithm 5.

### Open Targets retrieval

Open Targets exposes a GraphQL API rather than a relational schema. BioChirp therefore uses predefined retrieval functions instead of schema-graph planning for this database. The evaluated functions covered disease–target, drug–disease and gene–disease associations.

Entity mentions are resolved through the Open Targets search service. The highest-ranked canonical identifier is selected using a fixed procedure. Disease queries can additionally include descendant ontology terms. The resolved identifiers are inserted into predefined GraphQL operations rather than model-generated queries.

BioChirp retrieves successive pages until the endpoint reports that no further page is available or the configured operational limit is reached. Returned records are merged and deduplicated. Where required, records are ordered using predefined fields such as association score or clinical phase before conversion to the common BioChirp result format. The complete procedure is provided in Supplementary Algorithm 6.

### Natural-language presentation of retrieved evidence

After database execution, a language model can summarize the retrieved evidence. The model receives full-result statistics and a user-configurable preview of the returned rows. The default preview in the web interface contains 50 rows.

The model does not perform additional database searches and cannot modify the complete structured result. Changing the preview size affects only the generated summary.

### Evaluation design

BioChirp was evaluated at the component, retrieval-system and deployment levels (Fig. 2). Component experiments assessed multi-agent reasoning, schema mapping and entity grounding.

Retrieval benchmarks assessed count-level truncation, terminology robustness, repeated-run stability and coverage of curated database evidence. BioASQ-derived questions were used to compare BioChirp with agentic NL2SQL across multiple databases. Cross-database MCP workflows were evaluated qualitatively.

### Evaluating model coordination on MedQA

Model coordination was evaluated independently of database retrieval using 100 USMLE-style multiple-choice questions sampled from MedQA^29^. Each question had one accepted answer.

Four language models from different providers (Supplementary Fig. 3; Supplementary Table 11) answered every question independently and in parallel. Their responses were passed to a fifth model acting as the orchestrator. The orchestrator compared the candidate responses and selected the final answer, using web search only when needed to verify a disputed factual point.

We evaluated each standalone model, the coordinated system and the additional baseline configurations listed in Supplementary Table 12 over three independent runs. Accuracy was calculated as the proportion of questions for which the selected option matched the accepted MedQA answer.

### Evaluating biomedical encoders and language-model filtering

Entity grounding was evaluated using 43 biomedical terms comprising 14 diseases, 14 drugs and 15 genes. Diseases were sampled from ICD-11 categories, drugs from ATC categories and genes from Gene Ontology molecular-function categories (Fig. 5A–D). Curated Open Targets synonym sets served as the reference^20^.

Six biomedical sentence encoders (Supplementary Table 5) independently ranked the candidate terms for each query by cosine similarity. We compared two retrieval strategies in parallel. The fixed-k baseline retained the top k candidates, with k ranging from 25 to 725 in increments of 25. For Adaptive-S, the ranked cosine-similarity scores were examined for each query, and the point at which the scores began to drop sharply was used as the cutoff for retaining candidates. This knee point was identified using KneeLocator from the Python kneed package, with the sensitivity parameter set to S=5. For each encoder and retrieval strategy, the retrieved candidates were compared with the Open Targets synonym set for that query to calculate precision and recall.

Candidate retrieval was evaluated using precision and recall against the reference synonym sets. A retrieved term was counted as correct only when its text matched a reference synonym exactly after ignoring letter case.

Nine language models (Supplementary Table 6) were evaluated as zero-shot filters for candidate sets generated by three biomedical encoders (Supplementary Table 13). These encoders were selected because they showed balanced performance in the preceding encoder evaluation. All language models received the same filtering task and prompt (Fig. 6A). Precision and recall were measured before and after language-model filtering, and latency and inference cost were recorded for each model.

### Model Context Protocol retrieval benchmark

We compared twelve language-model and agentic configurations connected to the official Open Targets MCP endpoint (https://mcp.platform.opentargets.org/mcp) with the non-MCP BioChirp Open Targets pathway (Fig. 3A; Supplementary Table 1). The configurations and six benchmark questions are listed in Supplementary Table 1 and Supplementary Table 2. The experiment was designed as a stress test of exhaustive retrieval from large association sets rather than as an estimate of average MCP performance or biological answer accuracy.

The benchmark covered disease–target, drug–disease and gene–disease associations. Each association type included one non-exhaustive control question requesting top-ranked associations and one paired question requesting all matching associations. The top-ranked questions did not specify a fixed number of results because their purpose was to establish whether a configuration could perform the retrieval task without being required to exhaust the result set. They were not used to determine retrieval completeness. The paired exhaustive questions tested whether the same configuration could recover the complete association set. Each prompt was executed twice per configuration, giving twelve runs per configuration. BioChirp answered the same six prompts through its Open Targets retrieval pathway.

For each run, we recorded the number of associations returned, end-to-end latency and completion status. Outcomes were classified as no operational error, partial success, upstream retrieval failure, service-availability error, invocation timeout, exceeded interaction turns or malformed request. Detailed per-run outcomes are reported in Supplementary Table 14. BioChirp used exhaustive pagination within the configured operational limit for the exhaustive queries, and its reference outputs were manually inspected to confirm that they represented the requested association type.

Let *n_s,q,r_* denote the number of associations returned by system *s* for query *q* in run *r*. For the count-based comparison, we used the larger count observed across the two runs,

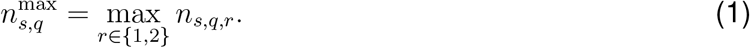

This best-case measure gives each MCP-connected configuration the benefit of its stronger run and avoids treating an isolated operational failure as evidence of retrieval incompleteness. Both individual runs are reported separately in Supplementary Table 14 so that run-to-run behaviour remains visible.

Zero counts were retained in all quantitative analyses. Count-based heatmaps used

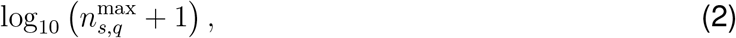

which permits visualization of zero counts without changing the underlying association counts.

For each exhaustive query *q*, let *n*_ref_*_,q_* denote the corresponding BioChirp reference count. Count yield was calculated as

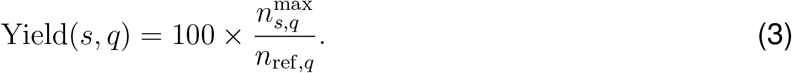

Count yield was interpreted as a measure of retrieval volume only for the three exhaustive queries. A count below the BioChirp reference establishes that the MCP-connected system did not return the complete association set. However, count yield does not measure record-level recall, and equal association counts do not establish that the two systems returned identical association identities. Because the benchmark questions were selected to stress exhaustive retrieval of large result sets, the experiment was designed to establish whether substantial truncation can occur under these conditions rather than to estimate its frequency across arbitrary MCP queries.

### NL2SQL robustness to biomedical synonyms

We evaluated whether agentic NL2SQL systems returned consistent results when the same biomedical entity was expressed using different accepted names. The benchmark contained 39 questions organized into 13 concept groups. Each group contained three versions of the same question that differed only in how the disease, drug or gene was named, using forms such as a preferred name, synonym, abbreviation, brand name or historical alias.

Twelve NL2SQL configurations combined four agentic frameworks with three language models. Each question was executed three times against the same HCDT snapshot, giving 1,404 NL2SQL runs. BioChirp was evaluated on the same questions and database snapshot. The questions and system configurations are listed in Supplementary Table 4 and Supplementary Table 3.

The NL2SQL prompt explicitly instructed each system to consider biomedical synonyms, abbreviations and aliases before generating SQL and to choose exact or substring matching according to the database field.

Let *n_s,g,v,r_* denote the number of rows returned by system *s* for concept group *g*, terminology variant *v* and run *r*. A concept group was classified as count-consistent for a system when

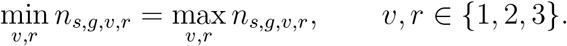

Thus, all three terminology variants had to return the same row count in all three runs. This measure assessed consistency of the number of retrieved records.

### Database field-mapping evaluation

Database field mapping was evaluated using 50 manually curated questions for HCDT and 50 for TTD. The questions covered different database columns, filter values, requested outputs and query structures. Each question was evaluated in three independent runs, producing 150 evaluations per database.

For each question, two language models independently identified which database columns should be used as filters, which columns should be returned and which values should be assigned to the filter columns. Both models used the same prompt and database information. Their outputs were evaluated separately and compared with the final coordinated field-mapping output. When the two models agreed, their common output was accepted. When they disagreed, an orchestrator reviewed both interpretations and produced the final field mapping.

An output was scored as correct only when all required database columns, their roles as filters or requested outputs, and the filter values assigned to those columns matched the manually defined reference. It was scored as incorrect if a required column was missing, a column was assigned the wrong role or a filter value was mapped to the wrong column.

### Repeated-query reproducibility and cross-system coverage

We compared three BioChirp backends with five language-model baselines (Supplementary Table 8). Each system answered the same 70 questions (Supplementary Table 9) across five independent runs under a fixed configuration. The questions covered drugs, targets, diseases and pathways.

BioChirp retrieved associations from Open Targets, TTD and HCDT. The language-model baselines answered using their internal model knowledge. Where identifiers were available, returned entities were normalized to canonical identifiers, including ChEMBL drug identifiers, Ensembl gene identifiers and disease ontology identifiers. Comparisons were based on identifier sets rather than surface wording, response order or repeated mentions. Reproducibility was calculated only for questions for which returned entities could be mapped to unique identifiers, preventing alternative names for the same entity from being counted as differences between runs.

Let *R_s,q,i_* denote the identifier set returned by system *s* for question *q* in run *i*. Reproducibility across the five runs was calculated as

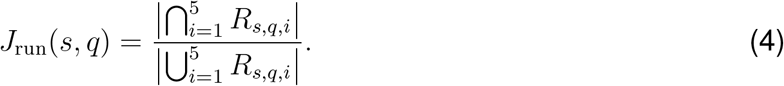

A value of one indicates that all five runs returned the same identifier set.

For cross-system comparisons, the identifier sets recovered across the five runs were first combined. Symmetric agreement between systems *s* and *s^′^* was then calculated using the Jaccard index,

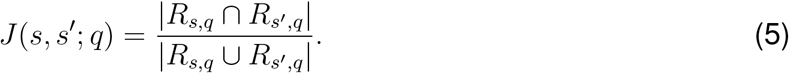

Directional coverage was calculated as

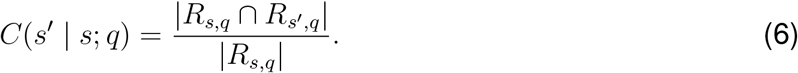

Here, *s* is the reference system, and the metric gives the fraction of its retrieved evidence also returned by *s^′^*. Directional coverage was calculated separately for each database. The heatmaps report the median metric value across all eligible questions.

HCDT and TTD were queried from local snapshots, whereas Open Targets was accessed through the live GraphQL API integrated into BioChirp. Open Targets reproducibility therefore refers to the database state during the evaluation period.

### BioASQ-derived database benchmark

We used BioASQ questions to compare BioChirp with agentic NL2SQL across multiple biomedical databases. The BioASQ source contained 5,729 unique factoid, yes/no, list and summary questions.

For each database, the complete BioASQ question set was reviewed together with the database schema using the ChatGPT web interface with GPT-5.6 Thinking. Questions that appeared answerable from the database were retained without changing their wording. A question could be assigned to more than one database.

This procedure produced 1,445 database–question evaluations from 1,013 unique questions across eleven databases. Open Targets contributed 535 evaluations but was excluded from the pairwise BioChirp–NL2SQL comparison because it uses GraphQL rather than a relational schema. The pairwise BioChirp–NL2SQL comparison therefore contained 910 database– question evaluations across ten relational databases. BioChirp and agentic NL2SQL received the same question set within each database.

The agentic NL2SQL baseline used LangChain with qwen-2.5-7b-instruct. CrewAI with Gemma-3-27B-IT and CrewAI with Qwen-2.5-7B-Instruct each returned the same row count across all terminology variants and repeated runs for 3 of 13 concept groups, compared with 2 of 13 for LangChain with Qwen-2.5-7B-Instruct. Because this difference was small and CrewAI required substantially longer runtimes in initial large-scale runs, LangChain was selected for the full BioASQ evaluation.

Retrieval and answer quality were evaluated separately. A table hit was recorded when a system retrieved at least one association from the assigned database. When BioChirp retrieved no association, its predefined fallback pathway generated a response using either language-model knowledge or web search. These cases were classified as having no table hit.

All responses were evaluated against the BioASQ reference using the PASS, PARTIAL and FAIL rubric. For the matched BioChirp–NL2SQL comparison, answer-quality statistics were calculated only for responses that retrieved at least one association from the assigned database (table hits). BioChirp responses classified as FAIL were then examined across the complete eleven-database evaluation to identify the causes of failure, including responses generated through the TABLE, LLM and WEB. PASS indicated that all expected reference elements were recovered. PARTIAL indicated that a subset of the expected elements was recovered. FAIL indicated that the expected answer was not produced. Associations retrieved from the assigned database but absent from the BioASQ reference were not marked as incorrect solely because they were absent from the reference.

Each response was evaluated three times by Claude Sonnet 5 using three predefined judge profiles for conservative factual assessment, biomedical domain-aware assessment and literal agreement with the reference. All profiles applied the same PASS, PARTIAL and FAIL definitions. A label selected by at least two judges became the final verdict. When the three judges returned different labels, a fourth Claude Sonnet 5 evaluation used a senior-adjudicator profile to select the verdict best supported by the retrieved evidence.

All BioChirp FAIL verdicts from the complete eleven-database evaluation were subsequently re-audited. Failures were assigned to database coverage gaps, genuine retrieval misses, incorrect final synthesis, incorrect routing or entity mapping, or borderline differences in evidence granularity. This audit included responses produced through table retrieval, language-model fallback and web fallback.

### BioChirp web-service implementation

BioChirp was deployed as a containerized microservice system with shared processing components and database-specific configurations.

Each local snapshot-backed database used a schema configuration describing its tables, columns, column descriptions and inferred relationships. A separate service configuration specified deployment parameters such as network port, worker count and memory allocation. Open Targets used a separate configuration because it was queried through a live GraphQL API.

Each database was exposed through a dedicated endpoint. Shared services handled database-field mapping, entity resolution and deterministic retrieval planning. The snapshot-backed databases used a common base prompt template, with database-specific instructions, constraints and examples added at runtime. For each incoming question, BioChirp also retrieved similar examples from a database-specific example collection and included them in the prompt. Qdrant stored vector representations of database column descriptions, biomedical entity names and example questions. Redis cached intermediate data, tracked query progress and temporarily stored results for access through shareable links.

### Prototype MCP interface for external agents

An MCP gateway exposed each BioChirp database endpoint as a tool that could be called by compatible external agents. The gateway was available at biochirp.iiitd.edu.in/mcp, with connection instructions at biochirp.iiitd.edu.in/connector. The gateway forwarded requests to the selected BioChirp database but did not choose which database to query or determine the sequence of database calls.

For the cross-database demonstration, we connected the gateway to Claude Desktop as the external MCP client. Claude first divided each composite question into a sequence of database-specific queries (Fig. 10). It then selected the database for each query and passed relevant entities retrieved at one step to subsequent queries. BioChirp processed each individual request through its standard retrieval workflow and returned the corresponding database records. Claude controlled the overall sequence of calls and generated the final synthesis from the retrieved evidence.

The MCP interface was used as a prototype to demonstrate cross-database access through an external agent. We did not systematically evaluate the accuracy, robustness or reproducibility of the cross-database orchestration performed through this interface. These workflows therefore demonstrate technical feasibility rather than quantitative performance. Completeness and reproducibility were evaluated only for individual BioChirp database retrieval calls, not for database selection, sequencing of calls or final synthesis by the external agent.

## Data availability

This study analysed publicly available third-party data from the Comparative Toxicogenomics Database^23^ (https://ctdbase.org/), the Highly Confident Drug-Target Database 2.0^27^ (http://hainmu-biobigdata.com/hcdt2/), the Open Targets Platform^20^ (https://platform.opentargets.org/), the Therapeutic Target Database^28^ (https://ttd.idrblab.cn/), the Human Phenotype Ontology^30^ (https://hpo.jax.org/), ClinVar^25^ (https://www.ncbi.nlm.nih.gov/clinvar/), Reactome^31^ (https://reactome.org/), the Molecular Signatures Database^32^ (https://www.gsea-msigdb.org/gsea/msigdb), Orphanet^24^ (https://www.orpha.net/), STRING^33^ (https://string-db.org/) and UniProt^34^ (https://www.uniprot.org/).

Ten resources were downloaded and converted into fixed local snapshots for BioChirp. These included HCDT 2.0, Reactome v96, STRING v12.0 and MSigDB v2026.1. Open Targets was accessed through its public GraphQL API on 27 February 2026 and was not stored as a local snapshot. Because Open Targets is a live resource, record counts obtained on later dates may differ from those reported here.

## Code availability

The BioChirp source code is available at https://github.com/abhi1238/biochirp under the MIT licence. The GitHub repository contains the application code, prompts, schema definitions, preprocessing workflows for the ten locally stored databases, the Open Targets API client, benchmark configurations, analysis scripts and documentation required to reproduce the reported evaluations.

A public deployment is available at https://biochirp.iiitd.edu.in/. The prototype Model Context Protocol server described in the Methods can be connected to an MCP-compatible client through https://biochirp.iiitd.edu.in/mcp, with setup instructions at https://biochirp. iiitd.edu.in/connector. The public service is under active development and may expose databases or interfaces beyond the 11-database configuration evaluated in this study.

## Acknowledgements

D.S. acknowledges support from the iHub-Anubhuti-IIITD Foundation under the NM-ICPS scheme of the Department of Science and Technology (DST), the Science and Engineering Research Board (SERB grant CRG/2022/007706), the Department of Biotechnology (DBT grant IC-12044(12)/4/2022-ICD-DBT), and the Indian Council of Medical Research (ICMR grant IIRPIG-2024-01-01321).

## Author contributions

D.S. conceived and supervised the study, contributed to figure preparation, and co-wrote, reviewed and edited the manuscript. A.H. developed the methodology, software and web server, designed the experiments, performed the experiments and analyses, conducted the literature review, deployed the system, prepared the figures, curated the GitHub repository and wrote the original manuscript. M.S. contributed to experimental design and analysis. R.K. contributed to the development of the Model Context Protocol interface. B.M., S.C.M.P. and P.M. validated the biological findings. B.M. also advised on database selection and assessed the design and operation of the tool from a biological perspective. N.B. contributed to figure preparation. O.C. and D.M. contributed to front-end development and deployed the web interface. G.A., M.F. and S.S. contributed to system development and deployed and hosted the backend infrastructure. P. and H.N. performed functional testing of the hosted web server. All authors reviewed and approved the final manuscript.

