## Supplementary Information for "Addressing challenges in agentic retrieval of structured data from biomedical databases"

Abhishek Halder [1,*], Manvendra Singh [2], Rohit Kesarwani [1,+], Bernadette Mathew [2,+], Namrata Bhattacharya [3,4], Om Chikhaliya [5], Dhruv Motwani [5], Sreeram Chandra Murthy Peela [2], Swarnava Samanta [2], Himanshu Nagdev [6], Piyush [7], Prateeksha Muddemmanavar [2], Mohammed Farooq [8], Gaurav Ahuja [2], and Debarka Sengupta [1,2,6,**]

[1]Department of Computer Science and Engineering, Indraprastha Institute of Information Technology Delhi, New Delhi 110020, India
[2]Department of Computational Biology, Indraprastha Institute of Information Technology Delhi, New Delhi 110020, India
[3]Australian Prostate Cancer Research Centre-Queensland, School of Biomedical Sciences, Faculty of Health, Queensland University of Technology, Brisbane, Queensland-4000, Australia
[4]Peter MacCallum Cancer Centre, Melbourne, VIC-3010, Australia
[5]HolboxAI, 701 Tillery Street Unit 12 1850, Austin, TX 78702
[6]Centre for Artificial Intelligence, Indraprastha Institute of Information Technology Delhi, New Delhi 110020, India
[7]MINDS NXT TECHNO VISION PVT LTD, Gulbarga, Karnataka 585101, India
[8]GeneSilico, Inc., 3267 Bee Caves Rd, STE 107-332, Austin, TX 78746, USA

[+]These authors contributed equally.

### SUPPLEMENTARY NOTES

#### Supplementary Note 1 — Model coordination improves biomedical reasoning outside structured database retrieval

Before evaluating coordination during structured database retrieval, we tested whether multiple language models could be combined to improve biomedical question answering. We hypothesized that models answering the same question independently would produce complementary interpretations and evidence. An orchestrator could then compare these responses, identify weaknesses and produce a more accurate final answer.

We sampled 100 USMLE-style multiple-choice questions from MedQA[29]. Each question had one expert-validated answer and required reasoning across clinical and biomedical concepts. We first measured the standalone accuracy of four language models from different providers (Supplementary Table 12).

We then sent each question to all four models in parallel. Their answers were passed to a fifth model acting as the orchestrator. The orchestrator compared the candidate responses and selected the final answer. When the candidate responses conflicted or a factual point required verification, the orchestrator could use web search to check that specific point before making the final selection (Supplementary Fig. 3A). Web search therefore served as a verification step within the coordination process rather than as an independent answer-generation pathway.

The coordinated system achieved accuracies of 95.0%, 95.0% and 94.0% across three runs, giving a mean accuracy of 94.7%. It outperformed every model evaluated independently. The model used as the orchestrator was the strongest standalone model and achieved 92.0% accuracy in each of the three runs. The coordinated system therefore achieved a mean accuracy 2.7 percentage points higher than the same model evaluated alone.

We also evaluated BioChatter, a model-agnostic biomedical LLM framework, using two underlying language models. Each BioChatter configuration closely reproduced the accuracy of its corresponding standalone model and provided no additional reasoning gain under the tested configuration.

These findings show that coordinating independently generated model responses can improve final answer selection in the implemented system. The orchestrator used web search only as an optional factual-verification step when resolving uncertainty or disagreement, rather than as an independent answer-generation pathway. The observed improvement therefore reflects the coordinated workflow as evaluated.

#### Supplementary Note 2 — Deterministic identifier assignment and schema assembly

BioChirp assigns one canonical identifier to each entity within each local database using a deterministic priority rule that selects the first available non-missing identifier. This procedure is applied independently within each database to support stable downstream joins, rather than to create a single identifier system shared across all sources.

When multiple identifier fields are available for the same entity, a fixed priority order is applied and the first non-missing value is selected. For example, pathway identifiers are resolved by checking available pathway-linked fields in order until a usable identifier is found, and gene identifiers are resolved in the same way from available gene-linked fields. Disease identifiers are assigned preferentially from structured disease-code fields; when no valid coded identifier is present, the disease name is used as a surrogate identifier within the local schema. This priority-based resolution is summarized in Supplementary Fig. 6.

For example, an HCDT pathway record can contain a placeholder pathway identifier alongside valid Reactome and KEGG identifiers. The placeholder pathway value is treated as missing, so the Reactome identifier is selected; KEGG is used only if Reactome is missing. For a TTD disease-linked record with no valid ICD-11 code, the cleaned disease name becomes the local disease identifier.

After identifier assignment, cleaned records are assembled into a fixed master–edge table schema (Supplementary Fig. 6). Master tables store one canonical identifier and one corresponding entity name per row, for example `drug_id` and `drug_name`. Association tables store paired identifiers representing relations between entity types, for example `drug_id` and `disease_id`. This design ensures that every relation resolves to valid entities and that downstream retrieval can be performed through deterministic, identifier-based joins.

Final validation removes null identifiers, duplicate relations and malformed or placeholder values before export. The resulting Parquet tables therefore provide a fixed retrieval schema with explicit entity nodes and reproducible relation edges.

### Supplementary Note 3 — Restricting schema mapping to queryable fields

Biomedical database schemas contain many columns that should not be treated as concepts in a natural-language question. BioChirp therefore defines a curated set of **queryable columns** for each database. These columns represent user-facing biomedical concepts, such as gene names, disease names, drug names and phenotype terms. Only queryable columns are considered when BioChirp determines which database fields should receive values from the question or which fields should be returned as the answer. Accordingly, user questions can query only these designated queryable fields, either as search constraints or as requested outputs.

This restriction improves mapping precision. Descriptive columns such as `synonyms`, `description` and `comment` contain useful supporting information, but exposing them to schema mapping can create incorrect field assignments and retrieval paths. For example, CTD contains a `synonyms` column in its disease, gene and chemical master tables. If these columns were treated as queryable concepts, a question such as "Which genes are associated with breast cancer?" could incorrectly connect the disease and gene tables through similarly named `synonyms` fields rather than through the gene–disease association table. BioChirp excludes such descriptive fields from schema mapping so that retrieval follows the relationships encoded in the database schema.

Non-queryable columns remain available to later stages when required. Internal identifier columns, such as `*_id`, provide the keys used by the deterministic planner and executor to join tables but are hidden from language-model field assignment because they are not normally expressed in user questions. Descriptive payload fields are also excluded from concept mapping, although they can be returned as supporting information when relevant. This separation limits language-model interpretation to user-facing biomedical concepts while preserving the complete relational structure needed for deterministic retrieval. The complete list of queryable fields defined for each supported database is available in the BioChirp GitHub repository (`https://github.com/abhi1238/biochirp`).

### Supplementary Note 4 — Matching alternative biomedical names using reference resources

BioChirp first checks whether a user-supplied drug, gene or disease term can be mapped using synonyms and identifiers obtained from biomedical databases, their official APIs or established biomedical reference resources. These mappings are used to resolve alternative names before lexical or semantic candidate generation.

The resources used depend on the entity type:

- **Drugs**: PubChem, RxNorm, ChEMBL, DrugNER and Open Targets, providing generic names, brand names, international non-proprietary names and other drug synonyms.
- **Genes**: HGNC, UniProt, MyGene.info, NCBI Gene and Open Targets, providing official gene symbols, previous symbols, aliases and alternative gene names.
- **Diseases**: NLM Clinical Tables, Disease Ontology, the Monarch disease terminology available through EBI OLS4 and Open Targets, providing preferred disease names and alternative disease terms.

For each query term, BioChirp normalizes the text by lowercasing and removing punctuation and compares it with the names and synonyms returned by the corresponding resources. A match is accepted when the normalized query term exactly matches a normalized reference name or synonym. The matched term is then mapped to the corresponding name or identifier used by the target database. Matches obtained at this stage are accepted without language-model validation.

**Worked example.** A user may query for *aspirin* when the target database represents the drug using *acetylsalicylic acid*. If a reference resource records *aspirin* as an alternative name for the same drug, BioChirp maps the user term to the corresponding database name or identifier and uses that value for retrieval without language-model validation.

### Supplementary Note 5 — Lexical matching for alternative entity names

Fuzzy lexical matching identifies database terms that are similar in spelling to the user-supplied term. This step is applied to unresolved disease names and to other free-text fields. It captures typographical errors, punctuation differences, abbreviated forms, changes in word order and partial word overlap.

BioChirp compares each query term only with values from the database column to which that term was assigned. For example, a disease term is compared only with disease names, not with drug or pathway names.

Four complementary string-similarity measures are evaluated:

1. **Global string similarity**: similarity across the complete query and candidate strings.
2. **Partial-string similarity**: similarity between the query and the most similar substring of the candidate, or vice versa.
3. **Reordered-token similarity**: word-level similarity after accounting for differences in token order.

4. **Shared-token similarity**: similarity based on the words shared by the query and candidate.

A candidate is retained when at least one similarity measure reaches the configured threshold of 90 on the 0–100 RapidFuzz scale. Candidates meeting this threshold are passed to language-model validation (Supplementary Note 7) before they can be used for retrieval.

**Worked example.** For the misspelled query term *tuberculsis*, fuzzy matching can identify the database term *tuberculosis* because of their high lexical similarity. The candidate is then passed to language-model validation before it is accepted.

### Supplementary Note 6 — Semantic matching for alternative entity names

Semantic retrieval provides a complementary candidate-generation route for disease names and other free-text fields. It is applied alongside lexical matching to recover alternative names with limited lexical overlap.

BioChirp encodes the query term using SapBERT[22] (Supplementary Table 5). The resulting vector is compared by cosine similarity with precomputed SapBERT representations of the names stored in the corresponding field of the target database.

The resulting candidates are ranked by cosine similarity. BioChirp uses the shape of the ranked similarity profile rather than a single fixed similarity threshold to determine how many candidates to retain. A knee point is identified at the sharpest decrease in similarity across the ranked candidates, providing a query-specific cutoff. Candidates above this cutoff are passed to language-model validation (Supplementary Note 7), whereas candidates below the cutoff are excluded.

### Supplementary Note 7 — Language-model validation of candidate entity names

Fuzzy and semantic retrieval are designed to recover a broad set of possible alternative names and can therefore introduce candidates that do not refer to the entity intended by the user. BioChirp uses a language model to remove these false-positive candidates before database retrieval.

**Model selection.** Gemma-4-31B was selected as the candidate-validation model because it provided balanced performance across the evaluated biomedical entity categories (Supplementary Table 6). It was therefore used to validate candidates generated by both fuzzy and semantic retrieval.

**Procedure.** The validator receives the original user-supplied term, the candidate names being evaluated and the entity type. It determines whether each candidate is a valid alternative name for the biomedical entity intended by the user. Each candidate receives an accept or reject decision. Accepted candidates are added to the resolved match set.

**Design rationale.** Fuzzy and semantic candidate generation favor recall and may therefore return false-positive matches. Language-model validation is used to restore precision before the resolved names are passed to deterministic database planning and retrieval.

### SUPPLEMENTARY METHODS

#### Supplementary Algorithm 1 — End-to-end BioChirp retrieval workflow

1. **Select the database:** Select the database associated with the chosen endpoint. When BioChirp is accessed through MCP, the external agent selects the appropriate database-specific BioChirp tool.
2. **Interpret the question:** identify the biomedical entities, the database fields that should constrain the search and the fields that should be returned.
3. **Resolve entities:** map user-supplied terms to the names or identifiers used by the selected database.
4. **Plan retrieval:** identify and connect the required tables for a relational database or select the corresponding predefined Open Targets operation.
5. **Retrieve records:** apply the finalized filters, perform the required joins or API calls, retrieve all matching records within the configured operational limits and remove duplicate records.
6. **Return the output:** provide the structured records together with the applied filters, retrieval path and total result count. Generate a readable summary from the result statistics and a bounded preview of the retrieved records.

#### Supplementary Algorithm 2 — Preprocessing a local biomedical database

1. Read the source tables and retain the identifier, name and association fields required by the database schema.
2. Standardize column names and values; remove source-specific prefixes, placeholders and malformed entries.
3. Split multi-valued cells so that each output row represents one entity or one relationship.
4. Assign one canonical identifier per entity using the fixed priority rule.
5. Construct master tables for entities and association tables for relationships.
6. Remove rows with missing identifiers and deduplicate the completed tables.
7. Write the fixed local snapshot and validate its schema before it is made available to the retrieval engine.

#### Supplementary Algorithm 3 — Language-model coordination for database field mapping

1. Rewrite the question independently with two language models to make the filter values and requested outputs explicit.
2. Embed each rewritten question using BAAI/bge-small-en-v1.5.
3. Retrieve the most similar queryable columns for each rewrite from the target database's schema index.
4. Use a language model to remove columns that are not relevant to each rewritten question.
5. Combine the retained columns from both rewrites into a single candidate schema.
6. Provide the same candidate schema to two independent language models and map each filter value and requested output to a database column.
7. Compare the database-column mappings produced by the two models programmatically.
8. Accept the mappings directly when both models agree. Otherwise, pass both interpretations to the orchestrator to produce the final mappings.
9. Pass the finalized mappings to entity resolution and retrieval planning.

#### Supplementary Algorithm 4 — Mapping biomedical terms to database vocabulary

1. Identify the type of each filter value and select the corresponding resolution procedure.
2. Match database identifiers, predefined coded values and standardized chemical representations directly to values stored in the database.
3. For gene and drug fields, map alternative names using synonyms and identifiers obtained from the corresponding database, its official API or established biomedical resources.
4. For disease names, combine these synonym sources with lexical matching and SapBERT-based semantic retrieval. For other free-text fields, generate candidates using lexical and semantic retrieval.
5. Evaluate candidates obtained by lexical or semantic retrieval with a language model and remove candidates that do not refer to the intended biomedical entity.
6. Combine directly matched names with validated candidates, map them to the corresponding database names or identifiers and remove duplicates.
7. Use the resolved names or identifiers as the final filter values for database planning and retrieval.

#### Supplementary Algorithm 5 — Retrieving records from relational databases

1. Validate the database snapshot, finalized filters, requested output fields and ordered retrieval plan.
2. Apply the finalized filters and traverse the selected foreign-key joins in the fixed plan order, retrieving all matching records within the configured operational limits.
3. Return the requested fields, remove duplicate records and compute statistics over the complete retrieved result.
4. Return the structured records together with the applied filters, selected tables, join path and full-result statistics. Provide a bounded preview of the records for language-model summarization.

#### Supplementary Algorithm 6 — Open Targets API retrieval

1. Route the interpreted request to a predefined Open Targets retrieval function.
2. Resolve the entity identifier and, for disease queries, add configured descendant terms.
3. Retrieve all available pages within the operational limit, then merge and deduplicate the records.
4. Rank and return the records in the common BioChirp result format.

### SUPPLEMENTARY FIGURES

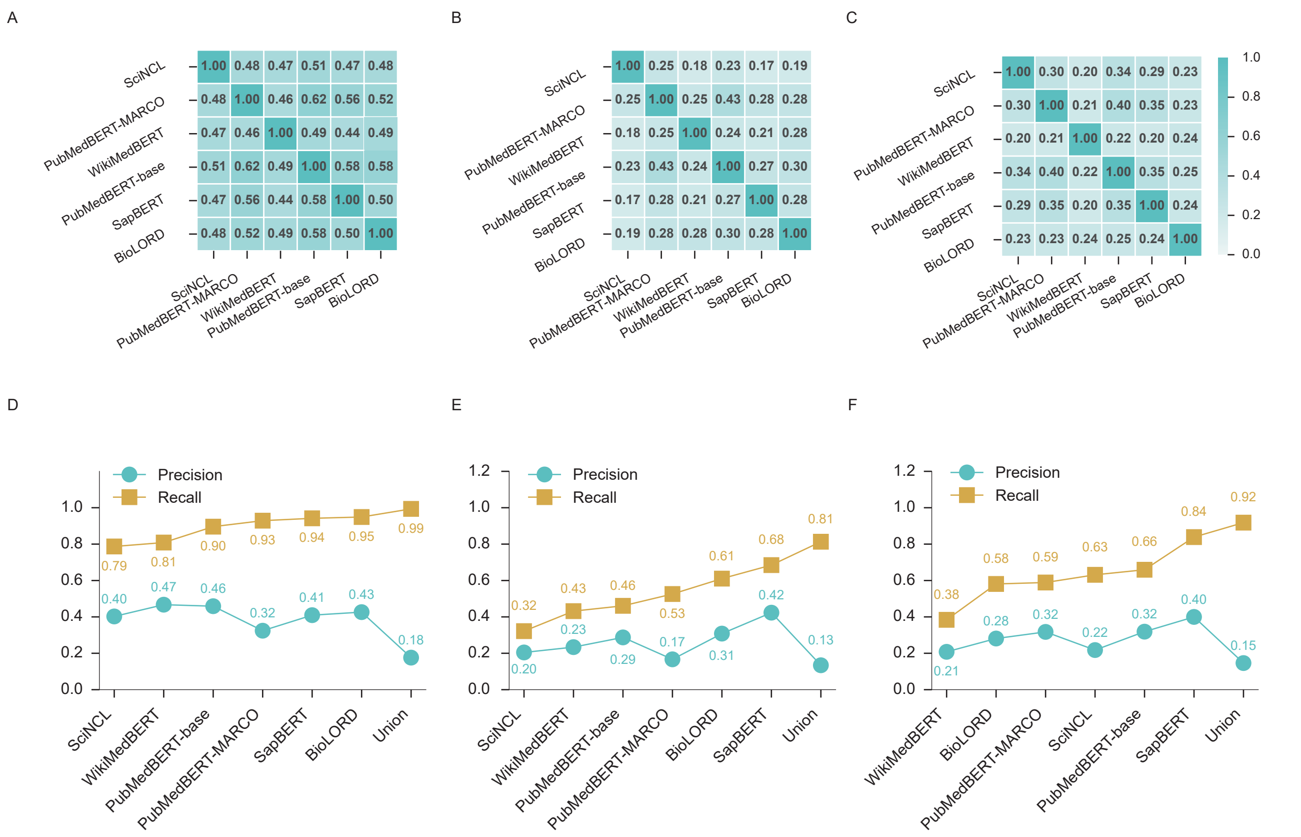


Supplementary Fig. 1: **Biomedical encoders retrieve complementary synonym candidate sets. A–C,** Pairwise Jaccard similarity between candidate sets returned at $k = 50$ by six biomedical sentence encoders for diseases, drugs and genes, respectively. Each cell reports the intersection divided by the union of the candidate sets returned by the corresponding encoder pair. The low overlap, particularly for drugs and genes, shows that encoder choice substantially changes which candidates reach downstream validation. **D–F,** Precision and recall of the six individual encoders and the union of their query-adaptive candidate sets for diseases, drugs and genes, respectively. Combining all encoders increased recall to 0.99 for diseases, 0.81 for drugs and 0.92 for genes, but reduced precision to 0.18, 0.13 and 0.15, respectively. The union therefore represents a recall-oriented strategy that transfers a larger false-positive burden to the downstream filter.

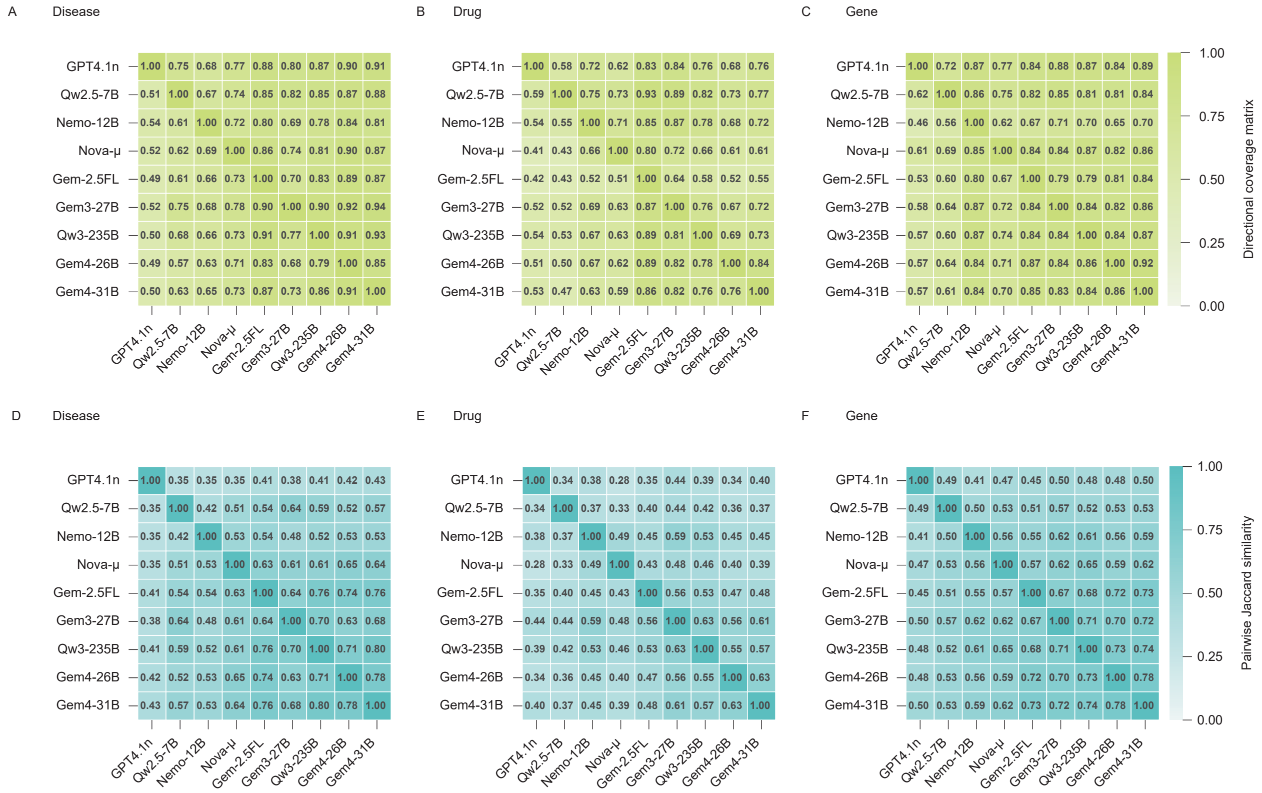


Supplementary Fig. 2: **Language-model filters differ in the breadth of synonym candidates they retain. A–C,** Directed coverage between candidate sets retained by nine language-model filters for diseases, drugs and genes, respectively. For row model $i$ and column model $j$, each cell reports $|S_i \cap S_j|/|S_i|$, the fraction of candidates retained by model $i$ that were also retained by model $j$. Because the denominator is the row model's set, the matrices distinguish conservative filters with small retained sets from broader filters that preserve more candidates. **D–F,** Symmetric pairwise Jaccard similarity between the same retained sets for diseases, drugs and genes, respectively. Agreement was lowest for drug candidates and highest for disease candidates, showing that model-specific filtering behaviour persisted even when all models evaluated identical candidate pools. [Model-label expansions: Nemo-12B (`mistralai/mistral-nemo`); Qw2.5-7B (`qwen/qwen-2.5-7b-instruct`); Qw3-235B (`qwen/qwen3-235b-a22b-2507`); Nova-$\mu$ (`amazon/nova-micro-v1`); Gem3-27B (`google/gemma-3-27b-it`); Gem4-26B (`google/gemma-4-26b-a4b-it`); Gem4-31B (`google/gemma-4-31b-it`); Gem-2.5FL (`google/gemini-2.5-flash-lite`); GPT4.1n (`openai/gpt-4.1-nano`).]

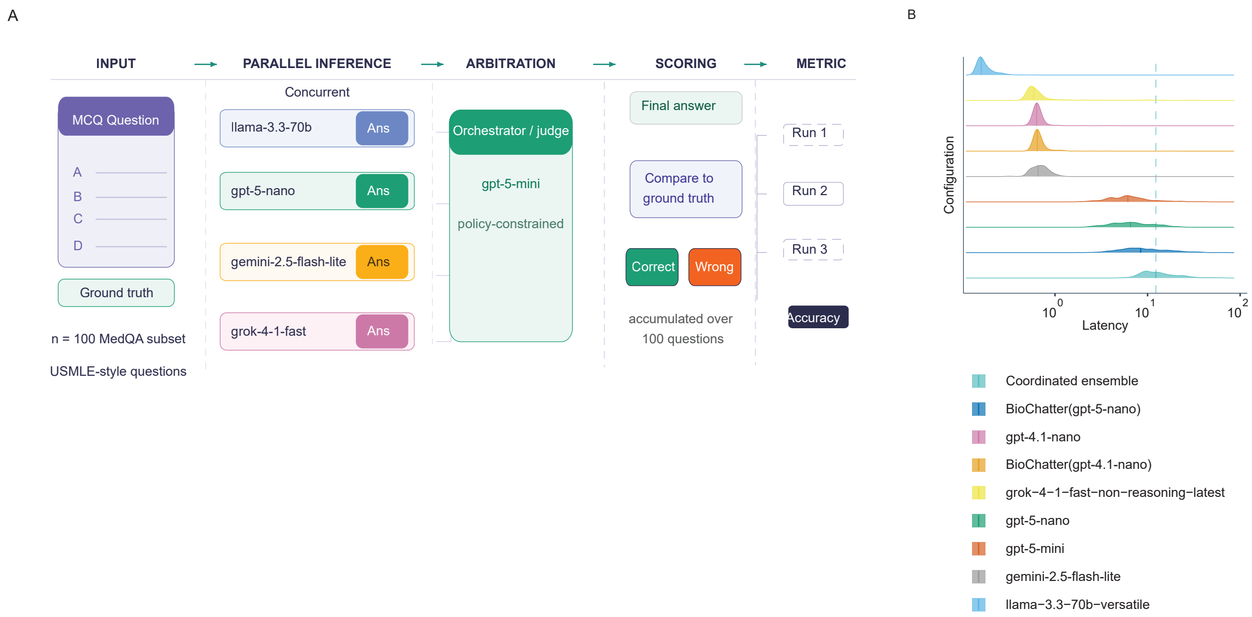


Supplementary Fig. 3: **Evaluation design and latency of judge-arbitrated multi-agent biomedical reasoning. A,** Evaluation workflow using 100 USMLE-style multiple-choice questions from MedQA. Llama-3.3-70B, GPT-5-nano, Gemini-2.5-Flash-Lite and Grok-4-1-Fast answered each question independently and concurrently. GPT-5-mini acted as a policy-constrained judge and produced one reconciled answer, which was compared with the expert-validated ground truth. Accuracy was calculated independently over three runs. **B,** Latency distributions across the 100-question evaluation for the coordinated ensemble, individual language models and BioChatter configurations, shown on a logarithmic scale. The coordinated ensemble incurred additional latency because it required parallel inference followed by judge arbitration. [Model-label expansions: Llama-3.3-70B (`llama-3.3-70b-versatile`); GPT-5-nano (`gpt-5-nano`); Gemini-2.5-Flash-Lite (`gemini-2.5-flash-lite`); Grok-4-1-Fast (`grok-4-1-fast-non-reasoning-latest`); GPT-5-mini (`gpt-5-mini`); GPT-4.1-nano (`gpt-4.1-nano`); BioChatter(gpt-5-nano) (`gpt-5-nano` through BioChatter); BioChatter(gpt-4.1-nano) (`gpt-4.1-nano` through BioChatter).]

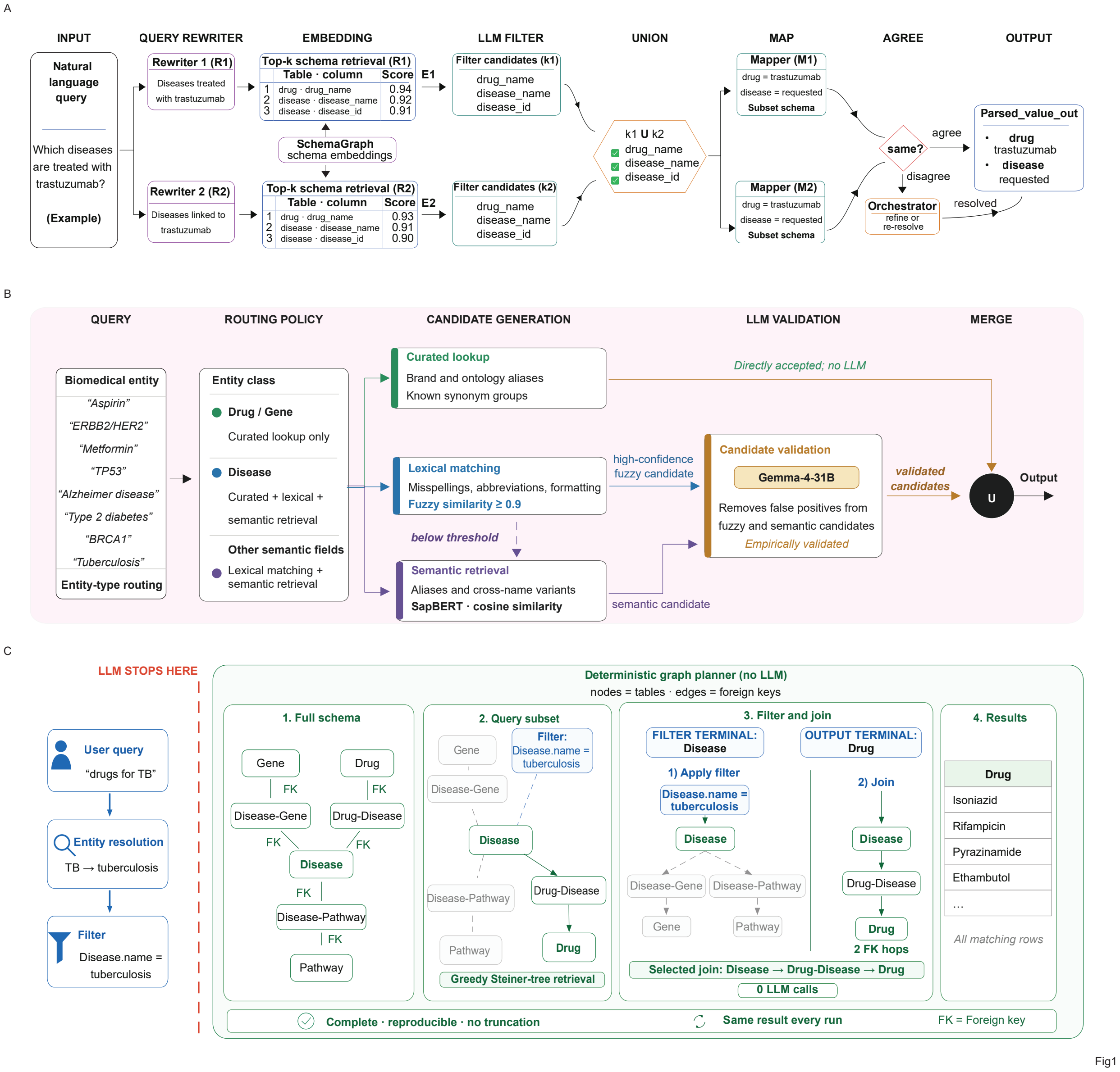


Supplementary Fig. 4: **Detailed BioChirp workflow for schema mapping, entity resolution and deterministic retrieval. A,** A natural-language query is independently rewritten, and the top-$k$ schema components most similar to each rewrite are retrieved from schema embeddings. This query-specific schema subset limits the context presented to the model, enabling large schemas to be handled without exposing the full schema. LLM-based filters retain relevant components, which are combined and mapped in parallel. Mapper agreement produces the parsed query representation, whereas disagreements are returned to the orchestrator for refinement or re-resolution. **B,** Field-specific entity-resolution policy. Drug and gene mentions are resolved using curated canonical names and synonym mappings. Disease names are resolved using curated synonyms, lexical matching and SapBERT-based semantic retrieval, whereas other free-text fields use lexical and semantic retrieval. Curated matches are accepted directly; lower-confidence lexical and semantic candidates are validated by Gemma-4-31B before the accepted candidates are merged. **C,** Deterministic schema-graph planning for the example query "drugs for TB". After TB is resolved to tuberculosis and mapped to the disease-name filter, language-model involvement ends. Tables are represented as nodes and inferred foreign-key relationships as edges. The planner identifies Disease and Drug as the filter and output terminals, respectively, and selects the minimal Disease–Drug-Disease–Drug path. The executor applies the filter, traverses the two validated foreign-key joins and returns all matching drug records for the fixed specification and database snapshot. [Model-label expansion: Gemma-4-31B (`google/gemma-4-31b-it`).]

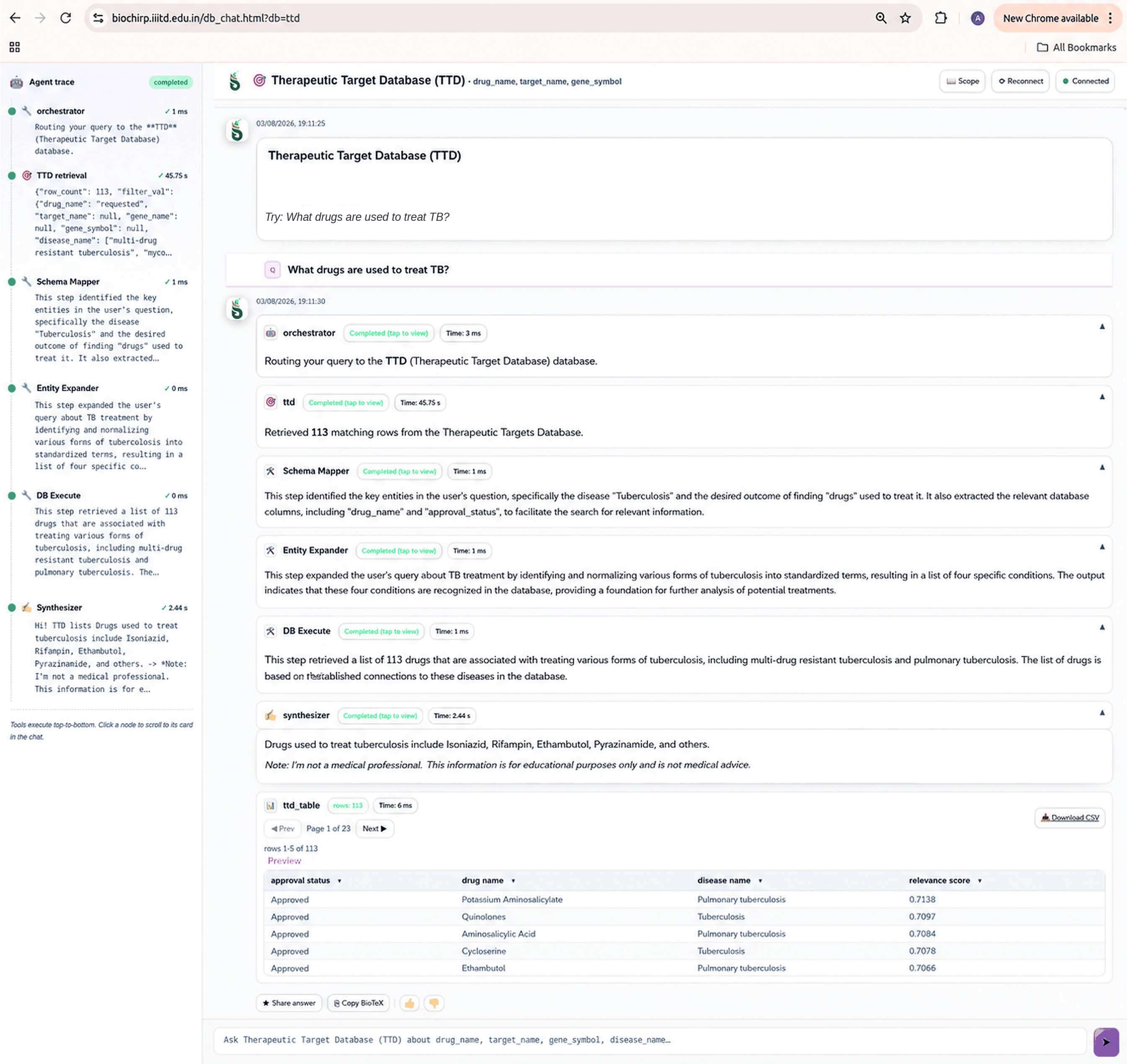


Supplementary Fig. 5: **BioChirp browser interface displays the execution trace and structured retrieval result.** Example query to the Therapeutic Target Database (TTD), "What drugs are used to treat TB?". The interface shows database routing, schema mapping, entity expansion, deterministic database execution and evidence-constrained synthesis as separate trace steps. Entity expansion mapped the query to four tuberculosis-related database terms, and execution retrieved matching drug records. The complete result is exposed through a paginated table and CSV download, while the natural-language response provides a concise summary of the retrieved evidence.

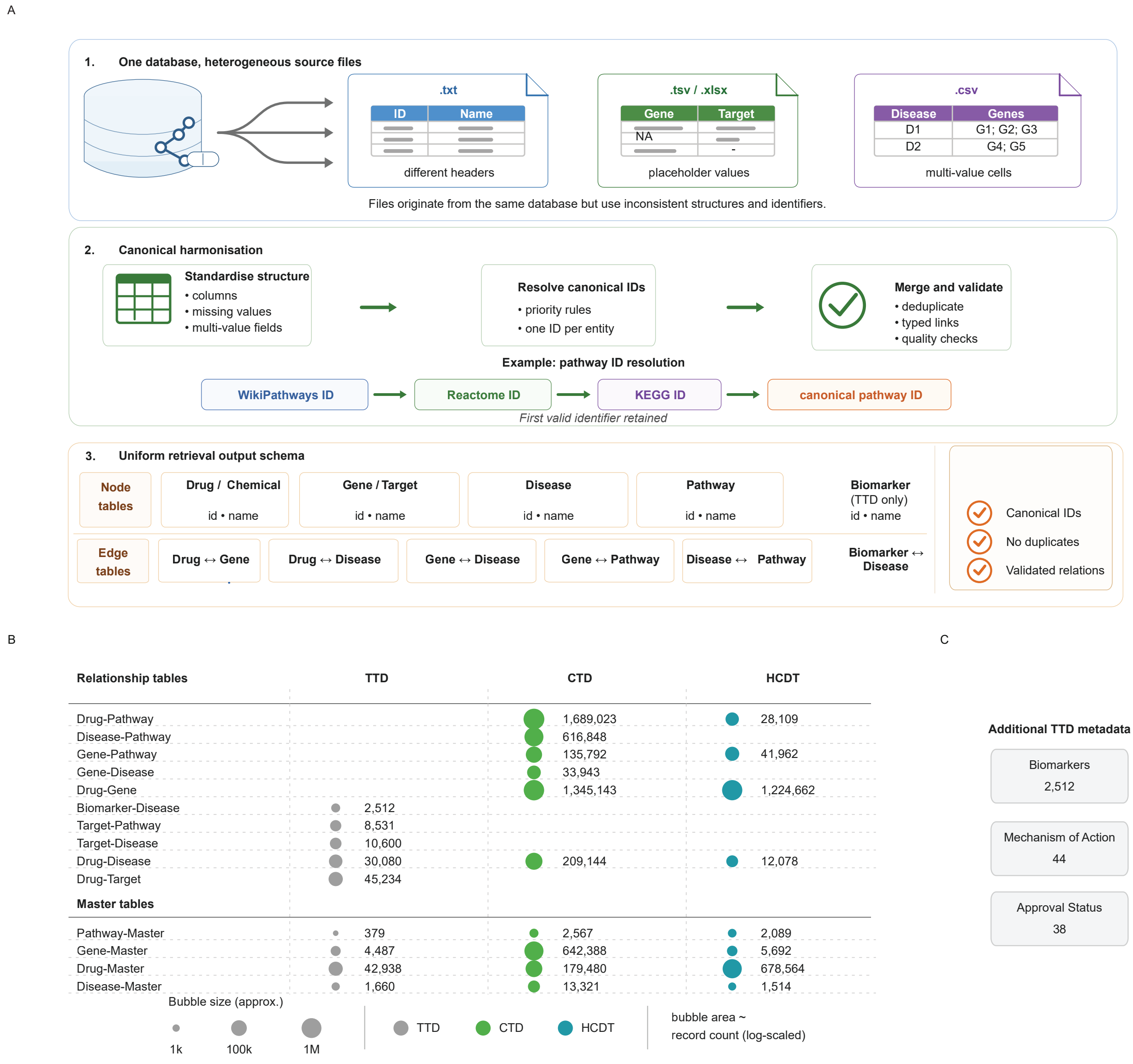


Supplementary Fig. 6: **Canonical preprocessing converts heterogeneous biomedical source files into validated relational snapshots.** **A,** Preprocessing workflow for the local TTD, CTD and HCDT databases. Source files from the same database can use different formats, headers, identifier systems, placeholder values and multi-valued cells. BioChirp standardizes columns and missing values, separates multi-valued records, assigns one canonical identifier per entity using deterministic priority rules, removes malformed or duplicate records and constructs a common retrieval schema comprising entity master tables and typed relationship tables. **B,** Numbers of records in the master and association tables produced for TTD, CTD and HCDT. For consistency across databases, the CTD chemical master table is shown under the harmonized label Drug-Master. The underlying CTD table remains a chemical master table. Bubble area represents the record count on an approximate logarithmic scale, with exact counts shown beside the corresponding bubbles. **C,** Additional TTD metadata retained after preprocessing, comprising 2,512 biomarker records, 44 mechanism-of-action categories and 38 approval-status categories. CTD, Comparative Toxicogenomics Database; HCDT, Highly Confident Drug–Target Database; TTD, Therapeutic Target Database.

### SUPPLEMENTARY TABLES

Supplementary Table 1: **Providers and models evaluated in the MCP-based retrieval benchmark.**

| Provider / environment | Model |
|---|---|
| OpenAI Agentic (Agents SDK) | gpt-4.1-mini |
| OpenAI Agentic (Agents SDK) | gpt-5-mini |
| OpenAI Agentic (Agents SDK) | gpt-5-nano |
| OpenAI Agentic (Agents SDK) | gpt-4o-mini |
| OpenAI Agentic (Agents SDK) | gpt-4.1-nano |
| OpenAI Chat Endpoint | gpt-5-nano |
| OpenAI Chat Endpoint | gpt-5-mini |
| OpenAI Chat Endpoint | gpt-4.1-mini |
| OpenAI Chat Endpoint | gpt-4.1-nano |
| OpenAI Chat Endpoint | gpt-4o-mini |
| Claude Desktop | Sonnet 4.5 |
| Claude Desktop | Haiku 3.5 |

Supplementary Table 2: **Query set used for the MCP retrieval benchmark.** Queries were designed to evaluate MCP-based biomedical retrieval across three association types and two retrieval modes.

| ID | Category | Mode | Query text |
|---|---|---|---|
| Q1 | Disease–target | Top-ranked | Provide the top-ranked molecular targets associated with chronic myeloid leukaemia (CML) in Open Targets. |
| Q2 | Disease–target | Exhaustive | Provide the complete list of molecular targets associated with chronic myeloid leukaemia (CML) in Open Targets. |
| Q3 | Drug–disease | Top-ranked | List the top diseases treated with aspirin according to Open Targets. |
| Q4 | Drug–disease | Exhaustive | Provide the complete list of diseases treated with aspirin according to Open Targets. |
| Q5 | Gene–disease | Exhaustive | Provide the complete list of diseases associated with the TP53 gene in Open Targets. |
| Q6 | Gene–disease | Top-ranked | List the top diseases associated with the TP53 gene in Open Targets. |

Supplementary Table 3: **Agentic frameworks and LLM models used in the agentic NL2SQL evaluation.**

| Framework | Model |
|---|---|
| PydanticAI | qwen/qwen-2.5-7b-instruct |
| PydanticAI | google/gemma-3-27b-it |
| PydanticAI | amazon/nova-micro-v1 |
| LangChain | qwen/qwen-2.5-7b-instruct |
| LangChain | google/gemma-3-27b-it |
| LangChain | amazon/nova-micro-v1 |
| CrewAI | qwen/qwen-2.5-7b-instruct |
| CrewAI | google/gemma-3-27b-it |
| CrewAI | amazon/nova-micro-v1 |
| PhiData | qwen/qwen-2.5-7b-instruct |
| PhiData | google/gemma-3-27b-it |
| PhiData | amazon/nova-micro-v1 |

Supplementary Table 4: Thirty-nine-query benchmark used for synonym-aware NL2SQL retrieval evaluation. Queries assigned to the same equivalent-query group are semantically matched and expected to return the same answer set.

| ID | Category | Equivalent group | Query text |
|---|---|---|---|
| Q1 | Disease (canonical) | Alzheimer treatment | Which drugs treat Alzheimer disease? |
| Q2 | Disease (possessive) | Alzheimer treatment | Which drugs treat Alzheimer’s disease? |
| Q3 | Disease (abbreviation) | Alzheimer treatment | Which drugs treat AD? |
| Q4 | Disease (canonical) | Parkinson treatment | Which drugs treat Parkinson disease? |
| Q5 | Disease (possessive) | Parkinson treatment | Which drugs treat Parkinson’s disease? |
| Q6 | Disease (abbreviation) | Parkinson treatment | Which drugs treat PD? |
| Q7 | Disease (canonical) | DLBCL treatment | What drugs treat diffuse large B-cell lymphoma? |
| Q8 | Disease (abbreviation) | DLBCL treatment | What drugs treat DLBCL? |
| Q9 | Disease (spacing variant) | DLBCL treatment | What drugs treat diffuse large B cell lymphoma? |
| Q10 | Gene (HGNC symbol) | ERBB2 targeting | Which drugs target ERBB2? |
| Q11 | Gene (clinical alias) | ERBB2 targeting | Which drugs target HER2? |
| Q12 | Gene (slash alias) | ERBB2 targeting | Which drugs target HER-2/neu? |
| Q13 | Gene (HGNC symbol) | KDR targeting | Which drugs target KDR? |
| Q14 | Gene (receptor alias) | KDR targeting | Which drugs target VEGFR-2? |
| Q15 | Gene (full enzyme name) | KDR targeting | Which drugs target vascular endothelial growth factor receptor 2? |
| Q16 | Gene (HGNC symbol) | FGFR2 targeting | Which drugs target FGFR2? |
| Q17 | Gene (retired symbol) | FGFR2 targeting | Which drugs target BEK? |
| Q18 | Gene (retired symbol) | FGFR2 targeting | Which drugs target KGFR? |
| Q19 | Drug (INN) | imatinib targets | Which genes does imatinib target? |
| Q20 | Drug (brand) | imatinib targets | Which genes does Gleevec target? |
| Q21 | Drug (development code) | imatinib targets | Which genes does STI571 target? |
| Q22 | Drug (INN) | sorafenib diseases | Which diseases is sorafenib approved for? |
| Q23 | Drug (salt form) | sorafenib diseases | Which diseases is sorafenib tosylate approved for? |
| Q24 | Drug (brand) | sorafenib diseases | Which diseases is Nexavar approved for? |
| Q25 | Drug (INN) | gefitinib targets | Which genes does gefitinib target? |
| Q26 | Drug (brand) | gefitinib targets | Which genes does Iressa target? |
| Q27 | Drug (development code) | gefitinib targets | Which genes does ZD1839 target? |
| Q28 | Gene (HGNC symbol) | EGFR targeting | Which drugs target EGFR? |
| Q29 | Gene (alias) | EGFR targeting | Which drugs target ErbB1? |
| Q30 | Gene (full name) | EGFR targeting | Which drugs target epidermal growth factor receptor? |
| Q31 | Gene (HGNC symbol) | ABL1 targeting | Which drugs target ABL1? |
| Q32 | Gene (fusion) | ABL1 targeting | Which drugs target BCR-ABL? |
| Q33 | Gene (c-prefix) | ABL1 targeting | Which drugs target c-ABL? |
| Q34 | Disease (canonical) | NSCLC treatment | Which drugs treat non-small cell lung cancer? |
| Q35 | Disease (hyphen variant) | NSCLC treatment | Which drugs treat non-small-cell lung cancer? |
| Q36 | Disease (abbreviation) | NSCLC treatment | Which drugs treat NSCLC? |
| Q37 | Disease (canonical) | myeloma treatment | Which drugs treat multiple myeloma? |
| Q38 | Disease (WHO term) | myeloma treatment | Which drugs treat plasma cell myeloma? |
| Q39 | Disease (abbreviation) | myeloma treatment | Which drugs treat MM? |

Supplementary Table 5: Biomedical sentence-embedding models evaluated for semantic candidate generation.

| # | Model identifier | Training and adaptation |
|---|---|---|
| 1 | `malteos/scincl` | SciBERT initialized; contrastively trained using neighbourhoods in the S2ORC citation graph. |
| 2 | `pritamdeka/S-PubMedBert-MS-MARCO` | PubMedBERT abstract-and-full-text model fine-tuned on MS MARCO using Sentence-Transformers. |
| 3 | `nuvocare/WikiMedical_sent_biobert` | BioBERT fine-tuned on medical Wikipedia sentence-similarity pairs. |
| 4 | `NeuML/pubmedbert-base-embeddings` | PubMedBERT fine-tuned on PubMed title–abstract pairs and similar-title pairs. |
| 5 | `cambridgeltl/SapBERT-from-PubMedBERT-fulltext-mean-token` | PubMedBERT adapted by self-alignment metric learning on English UMLS 2020AA synonyms. |
| 6 | `FremyCompany/BioLORD-2023` | `all-mpnet-base-v2` fine-tuned on ontology-derived BioLORD descriptions and LLM-generated AGCT definitions. |

Supplementary Table 6: **LLM models evaluated for biomedical synonym filtering via OpenRouter.** Models are ordered by output cost. All prices were verified on 2026- 06-01 via the OpenRouter API.

| # | Label | OpenRouter model slug | Provider(s) | $/1M in | $/1M out |
|---|---|---|---|---|---|
| 1 | nemo-12b | mistralai/mistral-nemo | DekaLLM, DeepInfra | 0.020 | 0.030 |
| 2 | qwen2.5-7b | qwen/qwen-2.5-7b-instruct | Phala | 0.040 | 0.100 |
| 3 | qwen3-235b | qwen/qwen3-235b-a22b-2507 | DeepInfra, WandB | 0.071 | 0.100 |
| 4 | nova-micro | amazon/nova-micro-v1 | Amazon Bedrock | 0.035 | 0.140 |
| 5 | gemma-3-27b | google/gemma-3-27b-it | DeepInfra, Novita | 0.080 | 0.160 |
| 6 | gemma-4-26b | google/gemma-4-26b-a4b-it | Cloudflare, DekaLLM, DeepInfra | 0.060 | 0.330 |
| 7 | gemma-4-31b | google/gemma-4-31b-it | DeepInfra, Novita, Parasail | 0.120 | 0.370 |
| 8 | gemini-2.5-flash-lite | google/gemini-2.5-flash-lite | Google AI Studio, Google | 0.100 | 0.400 |
| 9 | gpt-4.1-nano | openai/gpt-4.1-nano | OpenAI, Azure | 0.100 | 0.400 |

Supplementary Table 7: **Production entity-resolution filter model.**

| Model identifier | Provider | Function |
|---|---|---|
| gemma-4-31b | Google | Binary synonym-candidate filtering (fuzzy and semantic stages) |

Supplementary Table 8: Language-model systems evaluated as baselines against the deterministic database backends, with the exact model configuration used in each run.

| Model (exact identifier) | Provider |
|---|---|
| `grok-4-1-fast-non-reasoning-latest` | xAI |
| `llama-3.3-70b-versatile` | Groq |
| `gemini-2.5-flash-lite` | Google |
| `gpt-5-nano` | OpenAI |
| `gpt-5-nano` (LLM wrapper) | BioChatter (via OpenAI) |

Supplementary Table 9: **Benchmark question set used in the reproducibility and coverage evaluation.** Seventy natural-language queries spanning drug–disease, gene–disease, drug–target and gene–pathway association types. Each query was run five times per system.

| # | Question |
|---|---|
| 1 | Which diseases are associated with BRAF? |
| 2 | Which genes are associated with amyotrophic lateral sclerosis? |
| 3 | Which diseases are associated with CDK4? |
| 4 | Which pathways are associated with the JAK2 gene? |
| 5 | Which drugs are used to treat rheumatoid arthritis? |
| 6 | Which genes are associated with ovarian cancer? |
| 7 | What are the known targets of regorafenib? |
| 8 | Which pathways are associated with the STAT3 gene? |
| 9 | Which genes are associated with fever (pyrexia)? |
| 10 | Which diseases are treated with bevacizumab? |
| 11 | What are the known targets of cabozantinib? |
| 12 | Which pathways are associated with the TNF gene? |
| 13 | Which diseases are associated with the NPM1 gene? |
| 14 | Which pathways are associated with the CD19 gene? |
| 15 | Which drugs are used to treat migraine? |
| 16 | What drugs are used to treat rickets? |
| 17 | Which genes are associated with epilepsy? |
| 18 | Which diseases are treated with trastuzumab deruxtecan? |
| 19 | Which genes are associated with Alzheimer’s disease? |
| 20 | For which diseases is darolutamide used? |
| 21 | Which genes are associated with renal cell carcinoma? |
| 22 | Which diseases are associated with the KRAS gene? |
| 23 | Which pathways are associated with the APP gene? |
| 24 | Which genes are associated with schizophrenia? |
| 25 | Which diseases are treated with valdecoxib? |
| 26 | What are the known targets of vandetanib? |
| 27 | Which pathways are associated with the CFTR gene? |
| 28 | Which drugs are used to treat tuberculosis? |
| 29 | Which drugs are used to treat pancreatic adenocarcinoma? |

| # | Question |
|---|---|
| 30 | Which drugs are used to treat chronic myeloid leukaemia? |
| 31 | Which drugs are used to treat Parkinson's disease? |
| 32 | Which drugs are used to treat inflammatory bowel disease? |
| 33 | Which targets are acted on by imatinib? |
| 34 | Which biological targets does sorafenib interact with? |
| 35 | Which biological targets does tofacitinib interact with? |
| 36 | Which drugs target EGFR? |
| 37 | Which diseases are associated with TP53? |
| 38 | Which diseases are associated with FGFR2? |
| 39 | Which diseases are associated with HTT? |
| 40 | Which genes are associated with breast cancer? |
| 41 | Which genes are associated with colorectal cancer? |
| 42 | Which genes are associated with cystic fibrosis? |
| 43 | Which drugs are used to treat TNBC? |
| 44 | Which diseases are treated with metformin? |
| 45 | Which diseases are treated with adalimumab? |
| 46 | For which diseases is pembrolizumab used? |
| 47 | Which diseases are treated with sorafenib? |
| 48 | Which diseases are treated with nivolumab? |
| 49 | Which genes are associated with multiple sclerosis? |
| 50 | Which genes are associated with rheumatoid arthritis? |
| 51 | What are the known targets of abatacept? |
| 52 | Which genes are associated with lung cancer? |
| 53 | What are the known targets of ponatinib? |
| 54 | Which drugs are used to treat systemic lupus erythematosus? |
| 55 | Which genes are associated with autism spectrum disorder? |
| 56 | Which diseases are associated with the ROS1 gene? |
| 57 | Which diseases are associated with the ALK gene? |
| 58 | Which diseases are associated with the EGFR gene? |
| 59 | Which genes are associated with psoriasis? |
| 60 | Which genes are associated with ulcerative colitis? |
| 61 | Which genes are associated with tuberculosis? |
| 62 | Which genes are associated with hepatitis C? |
| 63 | Which genes are associated with Hodgkin lymphoma? |
| 64 | Which pathways are associated with TP53? |

| # | Question |
|---|---|
| 65 | Which drugs target ERBB2? |
| 66 | Which pathways are associated with the ADRB1 gene? |
| 67 | Which pathways are associated with the DRD2 gene? |
| 68 | Which pathways are associated with the HTR2A gene? |
| 69 | Which drugs target CD19? |
| 70 | Which drugs target CTLA4? |

Supplementary Table 10: **Benchmarking BioChirp on 4 curated MSigDB database questions.**

| Question | BioChirp | Official MSigDB Verdict with Claude Sonnet 5 as judge |
|---|---|---|
| What are the member genes of the human HALLMARK_APOPTOSIS gene set? | 161 genes, Homo sapiens | Official: "Further investigate these 161 genes" |
| Which human Hallmark gene sets contain both JUN and FOS? | 3 sets: TNFA_NFKB, P53_PATHWAY, HYPOXIA | Verified all 3 contain both JUN and FOS (200 genes each); APOPTOSIS has JUN but not FOS—correct exclusion |
| Which human Reactome pathways in MSigDB contain the gene BRCA2? | 13 rows, all Homo sapiens, R-HSA-* | Official: REACTOME_HOMOLOGY_DIRECTED_REPAIR → Homo sapiens, 140 genes, contains BRCA2; REACTOME_DNA_DOUBLE_STRAND_BREAK_REPAIR → 170 genes |
| What is the description of the HALLMARK_MYC_TARGETS_V1 gene set and how many genes does it contain? | 200 genes (human), "genes regulated by MYC" | Official: 200 genes, "A subgroup of genes regulated by MYC - version 1" |

Supplementary Table 11: **Language-model providers and model variants used in the question-answering benchmark.** The table lists the four language models used as the parallel question-answering agents.

| Provider | Model |
|---|---|
| OpenAI | gpt-5-nano |
| Google | gemini-2.5-flash-lite |
| xAI | grok-4-1-fast-non-reasoning-latest |
| Meta (via Groq) | llama-3.3-70b-versatile |

Supplementary Table 12: **Language-model providers, variants and wrapper configurations used in the MedQA benchmark.**

| Provider | Wrapper | Model |
|---|---|---|
| OpenAI | — | gpt-5-nano |
| OpenAI | — | gpt-5-mini |
| OpenAI | — | gpt-4.1-nano |
| Google | — | gemini-2.5-flash-lite |
| xAI | — | grok-4-1-fast-non-reasoning-latest |
| Meta (via Groq) | — | llama-3.3-70b-versatile |
| OpenAI | BioChatter | gpt-5-nano |
| OpenAI | BioChatter | gpt-4.1-nano |

Supplementary Table 13: **Fixed biomedical sentence encoders used to generate candidate pools for language-model filtering.** These three encoders generated the candidate sets evaluated by all nine zero-shot language-model filters.

| Encoder | Exact model identifier |
|---|---|
| SapBERT | `cambridgeltl/SapBERT-from-PubMedBERT-fulltext-mean-token` |
| BioLORD | `FremyCompany/BioLORD-2023` |
| PubMedBERT-MARCO | `pritamdeka/S-PubMedBert-MS-MARCO` |

Supplementary Table 14: **Detailed per-run benchmark results for MCP-connected systems and the non-MCP BioChirp comparator.** The table reports the query, model, run number, operational outcome, number of results returned, additional notes, latency and framework category for each benchmark run.

| Question category | Model | Run | Operational outcome | Count | Note | Latency (s) | Type |
|---|---|---|---|---|---|---|---|
| CML complete targets | gpt-4.1-nano | 1 | Upstream retrieval failure | 0 | — | 17.8 | Agentic |
| CML complete targets | gpt-4.1-nano | 2 | Upstream retrieval failure | 0 | GraphQL API Error | 18.7 | Agentic |
| CML complete targets | gpt-4o-mini | 1 | Exceeded interaction turns | 0 | — | 36.8 | Agentic |
| CML complete targets | gpt-4o-mini | 2 | Upstream retrieval failure | 0 | Open Targets Access Error | 35.8 | Agentic |
| CML complete targets | gpt-4.1-mini | 1 | Service-availability error | 0 | Open Targets API Unresponsive | 28.8 | Agentic |
| CML complete targets | gpt-4.1-mini | 2 | Exceeded interaction turns | 0 | — | 45 | Agentic |
| CML complete targets | gpt-5-nano | 1 | Exceeded interaction turns | 0 | — | 123.3 | Agentic |
| CML complete targets | gpt-5-nano | 2 | Partial success | 1 | — | 124.1 | Agentic |
| CML complete targets | gpt-5-mini | 1 | Malformed request | 0 | Your input exceeds the context window of this model. | 30.2 | Agentic |
| CML complete targets | gpt-5-mini | 2 | No operational error | 374 | — | 1192.3 | Agentic |
| CML top targets | gpt-4.1-nano | 1 | Partial success | 1 | — | 11.6 | Agentic |
| CML top targets | gpt-4.1-nano | 2 | Upstream retrieval failure | 0 | — | 10.9 | Agentic |
| CML top targets | gpt-4o-mini | 1 | Upstream retrieval failure | 0 | — | 28.3 | Agentic |
| CML top targets | gpt-4o-mini | 2 | Upstream retrieval failure | 0 | — | 17.1 | Agentic |
| CML top targets | gpt-4.1-mini | 1 | Exceeded interaction turns | 0 | — | 44.7 | Agentic |
| CML top targets | gpt-4.1-mini | 2 | Upstream retrieval failure | 0 | GraphQL API Error | 25.3 | Agentic |
| CML top targets | gpt-5-nano | 1 | Partial success | 1 | — | 123.1 | Agentic |
| CML top targets | gpt-5-nano | 2 | Partial success | 1 | — | 77.8 | Agentic |

#### Supplementary Table 14: Detailed per-run benchmark results for MCP-connected systems and the non-MCP BioChirp comparator (continued).

| Question category | Model | Run | Operational outcome | Count | Note | Latency (s) | Type |
|---|---|---|---|---|---|---|---|
| CML top targets | gpt-5-mini | 1 | Invocation timeout | 0 | — | 45.5 | Agentic |
| CML top targets | gpt-5-mini | 2 | Malformed request | 0 | Your input exceeds the context window of this model. | 44.8 | Agentic |
| Top diseases treated with aspirin | gpt-4.1-nano | 1 | Upstream retrieval failure | 0 | — | 14.6 | Agentic |
| Top diseases treated with aspirin | gpt-4.1-nano | 2 | Upstream retrieval failure | 0 | — | 7 | Agentic |
| Top diseases treated with aspirin | gpt-4o-mini | 1 | Upstream retrieval failure | 0 | — | 15.9 | Agentic |
| Top diseases treated with aspirin | gpt-4o-mini | 2 | No operational error | 10 | — | 34.3 | Agentic |
| Top diseases treated with aspirin | gpt-4.1-mini | 1 | No operational error | 29 | — | 23.5 | Agentic |
| Top diseases treated with aspirin | gpt-4.1-mini | 2 | Upstream retrieval failure | 0 | — | 28.8 | Agentic |
| Top diseases treated with aspirin | gpt-5-nano | 1 | No operational error | 12 | — | 81 | Agentic |
| Top diseases treated with aspirin | gpt-5-nano | 2 | Exceeded interaction turns | 0 | — | 124.3 | Agentic |
| Top diseases treated with aspirin | gpt-5-mini | 1 | No operational error | 168 | — | 259.4 | Agentic |
| Top diseases treated with aspirin | gpt-5-mini | 2 | No operational error | 168 | — | 286.2 | Agentic |
| All diseases treated with aspirin | gpt-4.1-nano | 1 | Upstream retrieval failure | 0 | — | 15.1 | Agentic |
| All diseases treated with aspirin | gpt-4.1-nano | 2 | Upstream retrieval failure | 0 | — | 11.3 | Agentic |
| All diseases treated with aspirin | gpt-4o-mini | 1 | Upstream retrieval failure | 0 | — | 33 | Agentic |
| All diseases treated with aspirin | gpt-4o-mini | 2 | Upstream retrieval failure | 0 | — | 12.8 | Agentic |
| All diseases treated with aspirin | gpt-4.1-mini | 1 | No operational error | 113 | — | 44 | Agentic |
| All diseases treated with aspirin | gpt-4.1-mini | 2 | No operational error | 168 | — | 68.7 | Agentic |

#### Supplementary Table 14: Detailed per-run benchmark results for MCP-connected systems and the non-MCP BioChirp comparator (continued).

| Question category | Model | Run | Operational outcome | Count | Note | Latency (s) | Type |
|---|---|---|---|---|---|---|---|
| All diseases treated with aspirin | gpt-5-nano | 1 | Invocation timeout | 0 | — | 68.4 | Agentic |
| All diseases treated with aspirin | gpt-5-nano | 2 | No operational error | 120 | — | 92.3 | Agentic |
| All diseases treated with aspirin | gpt-5-mini | 1 | No operational error | 168 | — | 955.7 | Agentic |
| All diseases treated with aspirin | gpt-5-mini | 2 | No operational error | 168 | — | 343.6 | Agentic |
| Top diseases associated with TP53 | gpt-4.1-nano | 1 | Upstream retrieval failure | 0 | — | 19.1 | Agentic |
| Top diseases associated with TP53 | gpt-4.1-nano | 2 | Upstream retrieval failure | 0 | — | 11.2 | Agentic |
| Top diseases associated with TP53 | gpt-4o-mini | 1 | Exceeded interaction turns | 0 | — | 53.5 | Agentic |
| Top diseases associated with TP53 | gpt-4o-mini | 2 | Upstream retrieval failure | 0 | — | 31.5 | Agentic |
| Top diseases associated with TP53 | gpt-4.1-mini | 1 | No operational error | 10 | — | 34.5 | Agentic |
| Top diseases associated with TP53 | gpt-4.1-mini | 2 | Upstream retrieval failure | 0 | — | 16.9 | Agentic |
| Top diseases associated with TP53 | gpt-5-nano | 1 | Exceeded interaction turns | 0 | — | 107 | Agentic |
| Top diseases associated with TP53 | gpt-5-nano | 2 | Exceeded interaction turns | 0 | — | 110.4 | Agentic |
| Top diseases associated with TP53 | gpt-5-mini | 1 | Malformed request | 0 | Your input exceeds the context window of this model. | 36.9 | Agentic |
| Top diseases associated with TP53 | gpt-5-mini | 2 | Malformed request | 0 | Your input exceeds the context window of this model. | 32.7 | Agentic |
| All diseases associated with TP53 | gpt-4.1-nano | 1 | Partial success | 1 | — | 7.3 | Agentic |
| All diseases associated with TP53 | gpt-4.1-nano | 2 | Partial success | 1 | — | 13.7 | Agentic |
| All diseases associated with TP53 | gpt-4o-mini | 1 | No operational error | 25 | — | 22.2 | Agentic |
| All diseases associated with TP53 | gpt-4o-mini | 2 | Exceeded interaction turns | 0 | — | 36.3 | Agentic |

#### Supplementary Table 14: Detailed per-run benchmark results for MCP-connected systems and the non-MCP BioChirp comparator (continued).

| Question category | Model | Run | Operational outcome | Count | Note | Latency (s) | Type |
|---|---|---|---|---|---|---|---|
| All diseases associated with TP53 | gpt-4.1-mini | 1 | No operational error | 46 | — | 40.3 | Agentic |
| All diseases associated with TP53 | gpt-4.1-mini | 2 | Upstream retrieval failure | 0 | — | 15.8 | Agentic |
| All diseases associated with TP53 | gpt-5-nano | 1 | No operational error | 10 | — | 128.9 | Agentic |
| All diseases associated with TP53 | gpt-5-nano | 2 | Upstream retrieval failure | 0 | — | 70.3 | Agentic |
| All diseases associated with TP53 | gpt-5-mini | 1 | Invocation timeout | 0 | — | 47.4 | Agentic |
| All diseases associated with TP53 | gpt-5-mini | 2 | Malformed request | 0 | — | 37.4 | Agentic |
| CML complete targets | gpt-4.1-nano | 1 | Upstream retrieval failure | 0 | — | 5.6 | ChatEndPoint |
| CML complete targets | gpt-4.1-nano | 2 | Upstream retrieval failure | 0 | — | 7.3 | ChatEndPoint |
| CML complete targets | gpt-4o-mini | 1 | Upstream retrieval failure | 0 | — | 13.4 | ChatEndPoint |
| CML complete targets | gpt-4o-mini | 2 | Upstream retrieval failure | 0 | — | 11.3 | ChatEndPoint |
| CML complete targets | gpt-4.1-mini | 1 | Upstream retrieval failure | 0 | — | 20.1 | ChatEndPoint |
| CML complete targets | gpt-4.1-mini | 2 | Upstream retrieval failure | 0 | — | 12.6 | ChatEndPoint |
| CML complete targets | gpt-5-nano | 1 | Malformed request | 0 | — | 53.5 | ChatEndPoint |
| CML complete targets | gpt-5-nano | 2 | No operational error | 10 | — | 130.6 | ChatEndPoint |
| CML complete targets | gpt-5-mini | 1 | Malformed request | 0 | — | 32.8 | ChatEndPoint |
| CML complete targets | gpt-5-mini | 2 | Malformed request | 0 | — | 32.6 | ChatEndPoint |
| CML top targets | gpt-4.1-nano | 1 | Upstream retrieval failure | 0 | — | 6.2 | ChatEndPoint |
| CML top targets | gpt-4.1-nano | 2 | Upstream retrieval failure | 1 | — | 5 | ChatEndPoint |

#### Supplementary Table 14: Detailed per-run benchmark results for MCP-connected systems and the non-MCP BioChirp comparator (continued).

| Question category | Model | Run | Operational outcome | Count | Note | Latency (s) | Type |
|---|---|---|---|---|---|---|---|
| CML top targets | gpt-4o-mini | 1 | Upstream retrieval failure | 0 | — | 7.2 | ChatEndPoint |
| CML top targets | gpt-4o-mini | 2 | Upstream retrieval failure | 0 | — | 8.2 | ChatEndPoint |
| CML top targets | gpt-4.1-mini | 1 | Upstream retrieval failure | 0 | — | 27.2 | ChatEndPoint |
| CML top targets | gpt-4.1-mini | 2 | Upstream retrieval failure | 0 | — | 11.8 | ChatEndPoint |
| CML top targets | gpt-5-nano | 1 | No operational error | 20 | — | 130.2 | ChatEndPoint |
| CML top targets | gpt-5-nano | 2 | Upstream retrieval failure | 0 | — | 40.7 | ChatEndPoint |
| CML top targets | gpt-5-mini | 1 | No operational error | 10 | — | 54.6 | ChatEndPoint |
| CML top targets | gpt-5-mini | 2 | Malformed request | 0 | — | 17.1 | ChatEndPoint |
| Top diseases treated with aspirin | gpt-4.1-nano | 1 | Upstream retrieval failure | 0 | — | 3.8 | ChatEndPoint |
| Top diseases treated with aspirin | gpt-4.1-nano | 2 | Upstream retrieval failure | 0 | — | 3.8 | ChatEndPoint |
| Top diseases treated with aspirin | gpt-4o-mini | 1 | Upstream retrieval failure | 0 | — | 10.2 | ChatEndPoint |
| Top diseases treated with aspirin | gpt-4o-mini | 2 | Upstream retrieval failure | 0 | — | 9.6 | ChatEndPoint |
| Top diseases treated with aspirin | gpt-4.1-mini | 1 | Upstream retrieval failure | 0 | — | 11.8 | ChatEndPoint |
| Top diseases treated with aspirin | gpt-4.1-mini | 2 | Upstream retrieval failure | 0 | — | 12.1 | ChatEndPoint |
| Top diseases treated with aspirin | gpt-5-nano | 1 | No operational error | 18 | — | 112.3 | ChatEndPoint |
| Top diseases treated with aspirin | gpt-5-nano | 2 | No operational error | 10 | — | 106.3 | ChatEndPoint |
| Top diseases treated with aspirin | gpt-5-mini | 1 | No operational error | 168 | — | 344.2 | ChatEndPoint |
| Top diseases treated with aspirin | gpt-5-mini | 2 | No operational error | 18 | — | 75.7 | ChatEndPoint |

#### Supplementary Table 14: Detailed per-run benchmark results for MCP-connected systems and the non-MCP BioChirp comparator (continued).

| Question category | Model | Run | Operational outcome | Count | Note | Latency (s) | Type |
|---|---|---|---|---|---|---|---|
| All diseases treated with aspirin | gpt-4.1-nano | 1 | Upstream retrieval failure | 0 | — | 6.2 | ChatEndPoint |
| All diseases treated with aspirin | gpt-4.1-nano | 2 | Upstream retrieval failure | 0 | — | 5.8 | ChatEndPoint |
| All diseases treated with aspirin | gpt-4o-mini | 1 | Upstream retrieval failure | 0 | — | 14.1 | ChatEndPoint |
| All diseases treated with aspirin | gpt-4o-mini | 2 | Upstream retrieval failure | 0 | — | 11.2 | ChatEndPoint |
| All diseases treated with aspirin | gpt-4.1-mini | 1 | Upstream retrieval failure | 0 | — | 10 | ChatEndPoint |
| All diseases treated with aspirin | gpt-4.1-mini | 2 | Upstream retrieval failure | 0 | — | 73.2 | ChatEndPoint |
| All diseases treated with aspirin | gpt-5-nano | 1 | No operational error | 68 | — | 89.5 | ChatEndPoint |
| All diseases treated with aspirin | gpt-5-nano | 2 | No operational error | 168 | — | 104.7 | ChatEndPoint |
| All diseases treated with aspirin | gpt-5-mini | 1 | No operational error | 168 | — | 75 | ChatEndPoint |
| All diseases treated with aspirin | gpt-5-mini | 2 | No operational error | 168 | — | 131 | ChatEndPoint |
| Top diseases associated with TP53 | gpt-4.1-nano | 1 | Upstream retrieval failure | 0 | — | 5 | ChatEndPoint |
| Top diseases associated with TP53 | gpt-4.1-nano | 2 | Upstream retrieval failure | 0 | — | 7.4 | ChatEndPoint |
| Top diseases associated with TP53 | gpt-4o-mini | 1 | Upstream retrieval failure | 0 | — | 7 | ChatEndPoint |
| Top diseases associated with TP53 | gpt-4o-mini | 2 | Upstream retrieval failure | 0 | — | 8.4 | ChatEndPoint |
| Top diseases associated with TP53 | gpt-4.1-mini | 1 | No operational error | 10 | — | 31 | ChatEndPoint |
| Top diseases associated with TP53 | gpt-4.1-mini | 2 | Upstream retrieval failure | 0 | — | 7.8 | ChatEndPoint |
| Top diseases associated with TP53 | gpt-5-nano | 1 | No operational error | 20 | — | 130.1 | ChatEndPoint |
| Top diseases associated with TP53 | gpt-5-nano | 2 | No operational error | 25 | — | 125.8 | ChatEndPoint |

#### Supplementary Table 14: Detailed per-run benchmark results for MCP-connected systems and the non-MCP BioChirp comparator (continued).

| Question category | Model | Run | Operational outcome | Count | Note | Latency (s) | Type |
|---|---|---|---|---|---|---|---|
| Top diseases associated with TP53 | gpt-5-mini | 1 | Malformed request | 0 | — | 19.8 | ChatEndPoint |
| Top diseases associated with TP53 | gpt-5-mini | 2 | Malformed request | 0 | — | 18.9 | ChatEndPoint |
| All diseases associated with TP53 | gpt-4.1-nano | 1 | Upstream retrieval failure | 0 | — | 8.8 | ChatEndPoint |
| All diseases associated with TP53 | gpt-4.1-nano | 2 | Partial success | 1 | — | 4.6 | ChatEndPoint |
| All diseases associated with TP53 | gpt-4o-mini | 1 | Upstream retrieval failure | 0 | — | 10.2 | ChatEndPoint |
| All diseases associated with TP53 | gpt-4o-mini | 2 | Partial success | 1 | — | 10.6 | ChatEndPoint |
| All diseases associated with TP53 | gpt-4.1-mini | 1 | Upstream retrieval failure | 0 | — | 9.8 | ChatEndPoint |
| All diseases associated with TP53 | gpt-4.1-mini | 2 | No operational error | 25 | — | 69.7 | ChatEndPoint |
| All diseases associated with TP53 | gpt-5-nano | 1 | Unclassified | 0 | — | 73.2 | ChatEndPoint |
| All diseases associated with TP53 | gpt-5-nano | 2 | No operational error | 20 | — | 123.6 | ChatEndPoint |
| All diseases associated with TP53 | gpt-5-mini | 1 | Malformed request | 0 | — | 15.4 | ChatEndPoint |
| All diseases associated with TP53 | gpt-5-mini | 2 | Malformed request | 0 | — | 20.6 | ChatEndPoint |
| CML top targets | Haiku 3.5 | 1 | Upstream retrieval failure | 0 | — | | Claude Desktop |
| Top diseases treated with aspirin | Haiku 3.5 | 1 | No operational error | 27 | — | | Claude Desktop |
| Top diseases associated with TP53 | Haiku 3.5 | 1 | No operational error | 5 | — | | Claude Desktop |
| CML complete targets | Haiku 3.5 | 1 | No operational error | 5 | — | | Claude Desktop |
| All diseases treated with aspirin | Haiku 3.5 | 1 | No operational error | 41 | — | | Claude Desktop |
| All diseases associated with TP53 | Haiku 3.5 | 1 | No operational error | 10 | — | | Claude Desktop |

#### Supplementary Table 14: Detailed per-run benchmark results for MCP-connected systems and the non-MCP BioChirp comparator (continued).

| Question category | Model | Run | Operational outcome | Count | Note | Latency (s) | Type |
|---|---|---|---|---|---|---|---|
| CML top targets | Haiku 3.5 | 2 | No operational error | 5 | — | | Claude Desktop |
| Top diseases treated with aspirin | Haiku 3.5 | 2 | Upstream retrieval failure | 0 | — | | Claude Desktop |
| Top diseases associated with TP53 | Haiku 3.5 | 2 | Upstream retrieval failure | 0 | — | | Claude Desktop |
| CML complete targets | Haiku 3.5 | 2 | No operational error | 5 | — | | Claude Desktop |
| All diseases treated with aspirin | Haiku 3.5 | 2 | No operational error | 36 | — | | Claude Desktop |
| All diseases associated with TP53 | Haiku 3.5 | 2 | No operational error | 20 | — | | Claude Desktop |
| CML top targets | Sonnet 4.5 | 1 | No operational error | 20 | — | | Claude Desktop |
| Top diseases treated with aspirin | Sonnet 4.5 | 1 | No operational error | 51 | — | | Claude Desktop |
| Top diseases associated with TP53 | Sonnet 4.5 | 1 | No operational error | 25 | — | | Claude Desktop |
| CML complete targets | Sonnet 4.5 | 1 | No operational error | 99 | — | | Claude Desktop |
| All diseases treated with aspirin | Sonnet 4.5 | 1 | No operational error | 99 | — | | Claude Desktop |
| All diseases associated with TP53 | Sonnet 4.5 | 1 | No operational error | 99 | — | | Claude Desktop |
| CML top targets | Sonnet 4.5 | 2 | No operational error | 10 | — | | Claude Desktop |
| Top diseases treated with aspirin | Sonnet 4.5 | 2 | No operational error | 55 | — | | Claude Desktop |
| Top diseases associated with TP53 | Sonnet 4.5 | 2 | No operational error | 20 | — | | Claude Desktop |
| CML complete targets | Sonnet 4.5 | 2 | No operational error | 99 | — | | Claude Desktop |
| All diseases treated with aspirin | Sonnet 4.5 | 2 | No operational error | 99 | — | | Claude Desktop |
| All diseases associated with TP53 | Sonnet 4.5 | 2 | No operational error | 99 | — | | Claude Desktop |

#### Supplementary Table 14: Detailed per-run benchmark results for MCP-connected systems and the non-MCP BioChirp comparator (continued).

| Question category | Model | Run | Operational outcome | Count | Note | Latency (s) | Type |
|---|---|---|---|---|---|---|---|
| CML top targets | BioChirp | 1 | No operational error | 4473 | — | 18.11 | BioChirp |
| Top diseases treated with aspirin | BioChirp | 1 | No operational error | 186 | — | 14.56 | BioChirp |
| Top diseases associated with TP53 | BioChirp | 1 | No operational error | 5631 | — | 16.81 | BioChirp |
| CML complete targets | BioChirp | 1 | No operational error | 4473 | — | 21.22 | BioChirp |
| All diseases treated with aspirin | BioChirp | 1 | No operational error | 186 | — | 14.5 | BioChirp |
| All diseases associated with TP53 | BioChirp | 1 | No operational error | 5631 | — | 14.11 | BioChirp |
| CML top targets | BioChirp | 2 | No operational error | 4473 | — | 15.58 | BioChirp |
| Top diseases treated with aspirin | BioChirp | 2 | No operational error | 186 | — | 12.76 | BioChirp |
| Top diseases associated with TP53 | BioChirp | 2 | No operational error | 5631 | — | 18.03 | BioChirp |
| CML complete targets | BioChirp | 2 | No operational error | 4473 | — | 17.82 | BioChirp |
| All diseases treated with aspirin | BioChirp | 2 | No operational error | 186 | — | 14.31 | BioChirp |
| All diseases associated with TP53 | BioChirp | 2 | No operational error | 5631 | — | 13.79 | BioChirp |
